# Imprinted antibody responses against SARS-CoV-2 Omicron sublineages

**DOI:** 10.1101/2022.05.08.491108

**Authors:** Young-Jun Park, Dora Pinto, Alexandra C. Walls, Zhuoming Liu, Anna De Marco, Fabio Benigni, Fabrizia Zatta, Chiara Silacci-Fregni, Jessica Bassi, Kaitlin R. Sprouse, Amin Addetia, John E. Bowen, Cameron Stewart, Martina Giurdanella, Christian Saliba, Barbara Guarino, Michael A. Schmid, Nicholas Franko, Jennifer Logue, Ha V. Dang, Kevin Hauser, Julia di Iulio, William Rivera, Gretja Schnell, Anushka Rajesh, Jiayi Zhou, Nisar Farhat, Hannah Kaiser, Martin Montiel-Ruiz, Julia Noack, Florian A. Lempp, Javier Janer, Rana Abdelnabi, Piet Maes, Paolo Ferrari, Alessandro Ceschi, Olivier Giannini, Guilherme Dias de Melo, Lauriane Kergoat, Hervé Bourhy, Johan Neyts, Leah Soriaga, Lisa A. Purcell, Gyorgy Snell, Sean P.J. Whelan, Antonio Lanzavecchia, Herbert W. Virgin, Luca Piccoli, Helen Chu, Matteo Samuele Pizzuto, Davide Corti, David Veesler

## Abstract

SARS-CoV-2 Omicron sublineages carry distinct spike mutations and represent an antigenic shift resulting in escape from antibodies induced by previous infection or vaccination. We show that hybrid immunity or vaccine boosters result in potent plasma neutralizing activity against Omicron BA.1 and BA.2 and that breakthrough infections, but not vaccination-only, induce neutralizing activity in the nasal mucosa. Consistent with immunological imprinting, most antibodies derived from memory B cells or plasma cells of Omicron breakthrough cases cross-react with the Wuhan-Hu-1, BA.1 and BA.2 receptor-binding domains whereas Omicron primary infections elicit B cells of narrow specificity. While most clinical antibodies have reduced neutralization of Omicron, we identified an ultrapotent pan-variant antibody, that is unaffected by any Omicron lineage spike mutations and is a strong candidate for clinical development.

---

The emergence of SARS-CoV-2 Omicron at the end of 2021 caused worldwide COVID-19 case surges. The Omicron BA.1 and BA.1.1 lineages swept the world followed by the BA.2 lineage (*1*). Although BA.1 and BA.2 share a large number of spike (S) mutations, they are each characterized by unique sets of amino acid changes, which are expected to be associated with different antigenic properties (*2, 3*). The BA.2.12.1 sublineage has recently emerged in the United States and is characterized by the presence of the S L452R (receptor-binding domain, RBD) and S704L (S_2_ subunit) mutations besides the BA.2-defining mutations. The BA.2.75 sublineage is spreading in multiple countries and carries unique mutations (added to the BA.2 background) in the N-terminal domain (NTD), the D339H, G446S and N460K mutations in the RBD along with the R493Q reversion (*4*). The BA.3 S glycoprotein comprises a combination of mutations found in BA.1 S and BA.2 S (*5*), whereas BA.4 S and BA.5 S comprise a deletion of residues 69-70, L452R and F486V and R493Q reversion compared to BA.2 S (*6*). We characterized the emergence of Omicron (BA.1) as a major antigenic shift due to the unprecedented magnitude of immune evasion associated with this variant of concern (*3, 7–11*). Mutations in the BA.1 S glycoprotein NTD and RBD, which are the main targets of neutralizing antibodies (*3, 12–17*), explain the markedly reduced plasma neutralizing activity of previously infected or vaccinated subjects, especially those that have not received booster doses, and escape from most monoclonal antibodies (mAbs) used in the clinic. As a result, an increasing number of reinfections or breakthrough infections are occurring (*18–20*), even though these cases tend to be milder than infections in immunologically naive individuals.

## Characterization of plasma and mucosal humoral responses to Omicron infection

Understanding the relationships between prior antigen exposure, through vaccination or infection with one SARS-CoV-2 strain, and the immune responses to subsequent infections with viruses from a different strain is paramount to guiding strategies to exit the COVID-19 pandemic. To address this question, we first benchmarked the magnitude of immune evasion associated with the BA.2 Omicron lineage by assessing the neutralizing activity of human plasma using a non-replicative vesicular stomatitis virus (VSV) pseudotyped with Wuhan-Hu-1 S harboring G614 (Wu-G614), Delta, BA.1, or BA.2 mutations or with SARS-CoV S (**Fig. 1A, Fig. S1A-F, Data S1**). We compared plasma from 6 cohorts of individuals: those previously infected in 2020 (with a Washington-1-like SARS-CoV-2 strain) and then vaccinated twice or three times (‘Infected-vaccinated 2 doses’, ‘Infected-vaccinated 3 doses’), those who were vaccinated and then experienced either a Delta or an Omicron BA.1 breakthrough infection (‘Delta breakthrough’ or ‘BA.1 breakthrough 2 doses’, ‘BA.1 breakthrough 3 doses’), or those that have only been vaccinated and boosted (‘vaccinated-only 3 doses’). Neutralizing antibody responses were slightly more robust against BA.2 than BA.1 among all groups except for the BA.1 breakthrough cases, with reductions of geometric mean titers (GMTs) relative to Wu-G614 ranging from 1.1-fold and 8.2-fold against BA.1 and between 1.8-fold and 4-fold against BA.2 (**Fig. 1A, Fig. S1A-F, Table S1**), in line with recent findings after primary vaccine series (*21*). All five cohorts experienced reductions in plasma neutralizing GMT of 1.5-3.6-fold against Delta (*22–24*) relative to Wu-G614, underscoring that even hybrid immunity (i.e., acquired through vaccination and infection (*25*)) do not overcome evasion from neutralizing antibody responses of this previously dominant variant of concern (**Fig. 1A, Fig. S1A-F, Table S1**). The highest neutralizing GMTs against SARS-CoV-2 variants were observed for BA.1 breakthrough cases, possibly due to exposure to BA.1 S, as no correlation was found between time intervals and GMTs **(Table S1)**. Neutralizing GMTs against the SARS-CoV S pseudovirus was reduced for all cohorts by 9 to 34-fold relative to Wu-G614, underscoring the marked genetic and antigenic divergence of this sarbecovirus clade (*18, 26, 27*).

**Figure 1.**
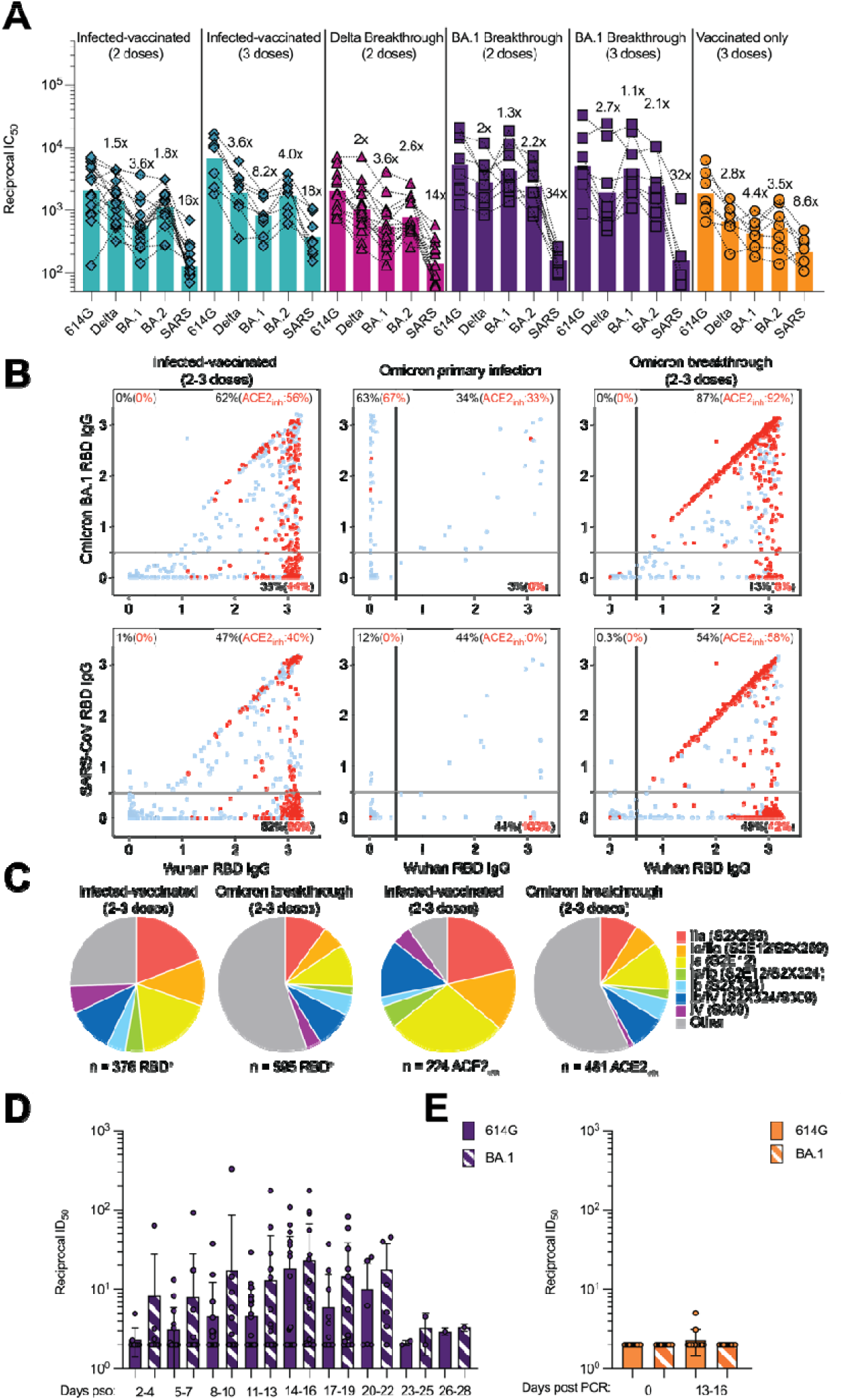
Evaluation of plasma, memory and mucosal antibody responses upon Omicron breakthrough infections in humans. **A**, Pairwise neutralizing activity (half-maximum inhibitory dose; ID_50_) against Wu-G614, Delta, BA.1, BA.2 and SARS-CoV S VSV pseudoviruses using plasma from subjects who were infected and vaccinated, vaccinated and experienced breakthrough infection, or vaccinated-only individuals. Vero E6-TMPRSS2 cell were used as target cells (*34*). Data are the geometric mean of an *n*lJ=lJ2 technical replicate and have been performed in at least 2 biologically independent experiments. GMTs are shown with a color-matched bar (and reported in Table S1) with fold change compared to Wu-G614 indicated above it. Demographics of enrolled donors are provided in Table S2. **B**, Cross-reactivity of IgGs secreted from memory B cells obtained from infected-vaccinated individuals (left, n=11), subjects who experienced a primary infection (center, n=3) or a breakthrough infection in January-March 2022 (right, n=7) when the prevalence of Omicron BA.1/BA.2 exceeded 90% in the region where samples were obtained (**Table S3**). Each dot represents a well containing oligoclonal B cell supernatant screened for the presence of IgGs binding to the SARS-CoV-2 Wuhan-Hu-1 and BA.1 RBDs (top) or to the SARS-CoV-2 Wuhan-Hu-1 and SARS-CoV RBDs (bottom) using ELISA. Red dots indicate inhibition of the interaction with ACE2 (using Wuhan- Hu-1 target antigen) as determined in a separate assay. The percentages are expressed relative to the total of positive hits against any of the antigen tested. Numbers of positive hits relative to individual donors are shown in Fig. S3. **C**, Frequency analysis of site-specific IgG antibodies derived from memory B cells. RBD sites targeted by IgG derived from memory B cells were defined by a blockade-of-binding assay using monoclonal antibodies specific for sites Ia (S2E12), Ib (S2X324), IIa (S2X259) and IV (S309). Lack of competition is indicated as “Other”. Pie charts show cumulative frequencies of IgGs specific for the different sites among total RBD-specific IgG antibodies (left) and those inhibiting binding of RBD to human ACE2 (right) in 11 infected-vaccinated individuals or 7 breakthrough cases. **D**, Neutralizing activity against Wu-G614 and BA.1 S VSV pseudoviruses in nasal swabs obtained longitudinally upon BA.1 breakthrough infection up to 28 days following symptom onset (pso). **E**, Neutralizing activity against Wu-G614 and BA.1 S VSV pseudoviruses in nasal swabs obtained longitudinally following a negative PCR test of vaccinated-only individuals.

Given the recall of Wuhan-Hu-1 plasma neutralizing antibodies in Omicron breakthrough cases, we investigated the cross-reactivity of RBD-specific antibodies produced by in vitro stimulated memory B cells or circulating plasma cells collected soon after infection (*28*). These analyses used blood samples from individuals who were infected prior to the emergence of Omicron and subsequently vaccinated as well as subjects who experienced primary Omicron infections or Omicron breakthrough infections. Primary and breakthrough infections occurred between January and March 2022 during which the prevalence of Omicron BA.1/BA.2 exceeded 90% in the region from which samples were obtained (**Fig. S2**). Strikingly, more than 80% of IgG antibodies secreted by memory B cells and plasma cells obtained from breakthrough cases cross-reacted with the Wuhan-Hu-1, BA.1, BA.2 and Delta RDBs, and more than 90% of these antibodies blocked binding to ACE2 (a correlate of neutralization (*12, 29*)) (**Fig. 1B-C, Fig. S3-S5, Table S2**). Furthermore, Omicron breakthrough infections failed to elicit BA.1- or BA.2 RBD-specific antibodies. These findings illustrate how immunological imprinting from prior exposure, i.e., ‘original antigenic sin’, can strongly affect the response to antigenically novel antigens. Conversely, memory B cell-derived RBD-specific IgG antibodies obtained from Omicron primary infections were present at low frequency and were mostly specific for BA.1 and BA.2, highlighting their antigenic relatedness (**Fig. 1B-D, Fig. S3-S6, Table S2**). The frequency of IgG antibodies cross-reacting with the SARS-CoV RBD was similar across all three cohorts, concurring with neutralization data with matched sera (**Table S3**). Plasma neutralizing activity of Omicron-infected (primary or breakthrough) cases was reduced on average 6.1-fold against BA.5 relative to BA.1 (**Table S3**), likely as a result of the L452R and F486V additional RBD mutations in the former lineage, concurring with recent data (*30*).

The site specificity of RBD-specific antibodies produced from stimulated memory B cells was determined by competition with structurally characterized mAbs targeting four distinct antigenic sites (*12*) (**Fig. S7**). Antibodies recognizing antigenic sites Ia and Ib which overlap with the receptor-binding motif (RBM), such as mAbs S2E12 (*31*), and S2X324, are found at lower frequency upon Omicron breakthrough infections relative to infected-vaccinated subjects, consistent with the presence of several immune escape mutations in the Omicron RBM (*3, 17*). A similar relative reduction was observed for antibodies targeting the RBD antigenic site IIa recognized by the S2X259 mAb (*32*), in agreement with previous findings on the Omicron immune escape from several site IIa mAbs (*3, 17*). Whereas most memory B cell-derived antibodies from (pre-Omicron) infected-vaccinated individuals competed with the four reference mAbs used, a large fraction of antibodies from Omicron breakthrough cases did not compete with any of these four mAbs, indicating they recognize other undefined RBD antigenic sites. Collectively, these findings demonstrate that infection with Omicron preferentially expanded existing B cell pools primed by vaccination and elicited cross-reactive plasma cells and antibodies, supporting the concept of immunological imprinting.

To evaluate mucosal antibody responses in subjects who experienced a BA.1 breakthrough infection or vaccinated-only individuals, we assessed IgG and IgA binding titers in nasal swabs. Although we detected S-specific IgG, and to a lesser extent IgA, in swabs from several breakthrough cases, vaccinated-only individuals had no detectable titers of binding antibodies **(Fig. S8)**. Furthermore, we measured mucosal neutralizing activity against G614 and BA.1 S VSV pseudoviruses for breakthrough samples up to a month post symptom onset **(Fig. 1D-E, Data S1)**. We note that the magnitude of neutralizing antibody responses in nasal swabs cannot be directly compared to plasma samples due to the self-administration procedure and resulting sample non-uniformity. Overall, we observed heterogenous polyclonal neutralizing antibody responses among BA.1 breakthrough cases but not in vaccinated-only individuals, in line with previous data indicating that mRNA vaccines or adenovirus-vectored vaccines do not induce appreciable mucosal antibody responses upon intra-muscular delivery (*33*).

## Omicron sublineages escape neutralization mediated by most clinical mAbs

We next evaluated the impact of BA.1, BA.2, BA.3, BA.4, BA.5, BA.2.12.1 and BA.2.75 S mutations on neutralization mediated by a panel of RBD-directed mAbs using VSV pseudoviruses and Vero E6 cells as target cells. The site Ia/Ib COV2-2130 mAb poorly neutralized BA.1 (*3*), while it neutralized BA.2, BA.3, BA.4, BA.5, BA.2.12.1 and BA.2.75 with 1.6-fold, 4.2-fold, 14.5-fold, 8.8-fold, 2.0-fold and 7.9-fold respective drops in half-maximal inhibition concentrations (IC_50_s) compared to Wu-G614. Moreover, the COV2-2196+COV2-2130 mAb cocktail experienced a 106.4-fold, 6.3-fold, 35-fold, 92.8-fold, 46.5-fold, 9.7-fold and 9.1-fold reductions in potency against BA.1, BA.2, BA.3, BA.4, BA.5, BA.2.12.1 and BA.2.75, respectively **(Fig. 2A)**. Since COV2-2196 poorly inhibited Omicron sublineages (except for BA.2.75 where the drop in IC_50_ is 17.3-fold), the neutralizing activity of the cocktail is largely mediated by COV2-2130. The only mutational difference within the COV2-2130 epitope is at position 446 which is a glycine residue in Wuhan-Hu-1, BA.2, BA.4 and BA.5 or a serine residue in BA.1, BA.3 and BA.2.75, the latter residue disrupting the binding interface of COV2-2130 (*17*). The importance of this site was also identified through deep mutational scanning (*35*) and this point mutation was shown to reduce neutralizing activity ∼4 fold for COV2-2130 (*7*). The greater reduction in potency against BA.4 and BA.5 relative to BA.2 and BA.3 is likely driven by the L452R mutation, as reported (https://www.fda.gov/media/154701/download) (*35*). The REGN10987+REGN10933 mAb cocktail, LY-CoV16+LY-CoV555 mAb cocktail, the CT-P59 mAb and ADI-58125 mAb experienced reductions of in vitro neutralization potency ranging between two and four orders of magnitude against all Omicron sublineages compared to G614 due to mutations in the RBM (*17*). CT-P59, however, retained neutralizing activity against the BA.2.75 sublineage (29.2-fold reduction relative to Wu-G614). The recently described ACE2-mimicking S2K146 mAb (*36*) that retained unaltered activity against BA.1 compared to Wu-G614 (*3*), had a mildly reduced neutralizing activity against BA.2, BA.3, BA.2.12.1 and BA.2.75 (3.3-fold, 3.1-fold, 1.9-fold and 4.3-fold, respectively). However, S2K146 displayed a marked reduction in neutralizing activity against BA.4 and BA.5 (472-fold and 285-fold drops in IC50s compared to Wu-G614), likely due to the F486V mutation.

**Figure 2:**
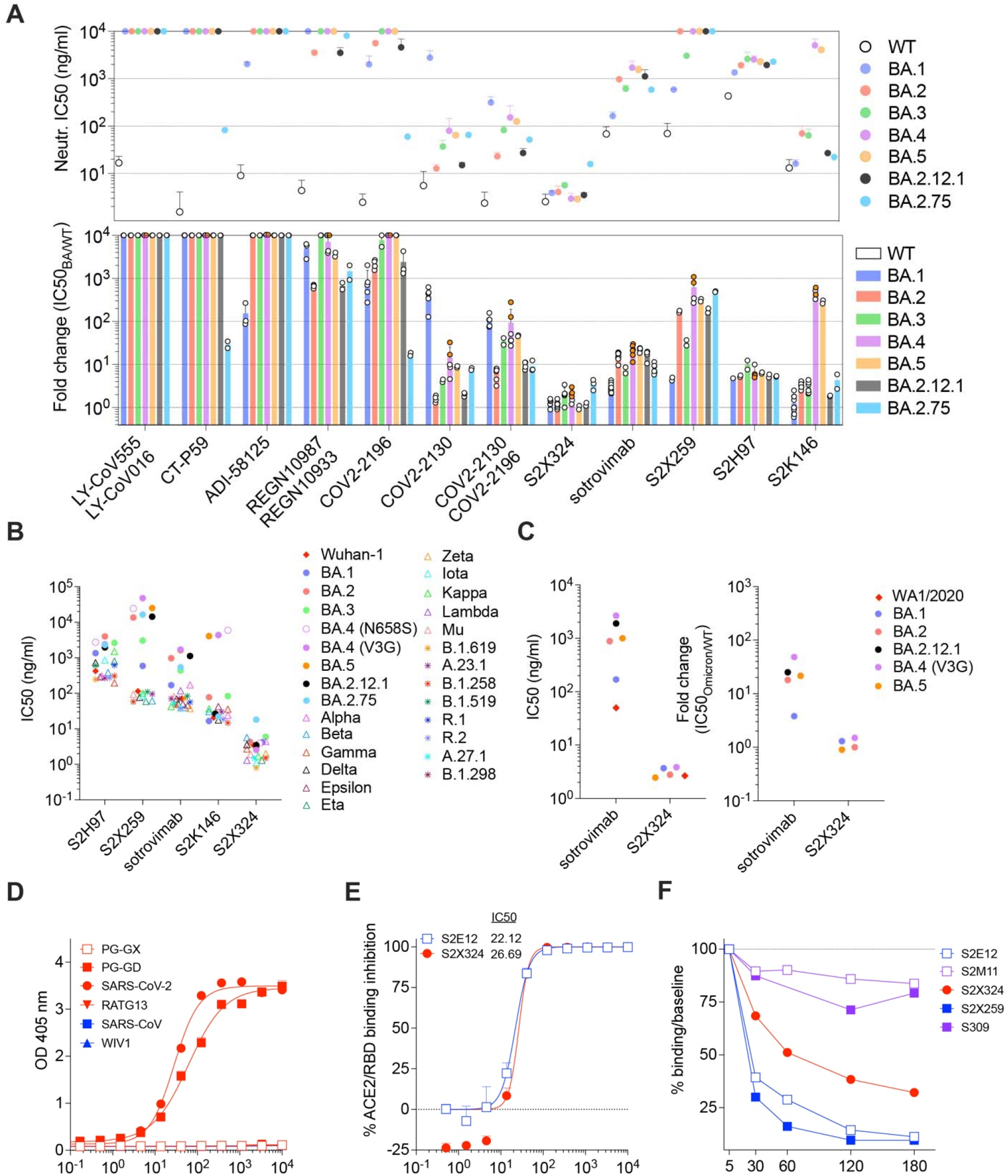
Identification and characterization of S2X324 as a pan-variant RBD-targeted mAb. **(A)** mAb-mediated neutralization of BA.1, BA.2, BA.3, BA.4, BA.5, BA.2.12.1 and BA.2.75 S VSV pseudoviruses. The potency of each mAb or mAb cocktail is represented by their IC_50_ (top, geometric mean ± SD) or fold change relative to neutralization of the Wuhan-Hu- 1 (D614) pseudovirus (bottom, average ± SD). Two haplotypes of BA.4 S were tested: BA.4-V3G (orange dots) and BA.4-N658S (white dots). **(B)** Neutralization of SARS-CoV-2 VSV pseudoviruses mediated by broadly neutralizing sarbecovirus mAbs. Each symbol represents the Geometric mean of IC_50_ values of at least two independent experiments. **(C)** Neutralization of SARS-CoV-2 authentic viruses by sotrovimab and S2X324 expressed as the geometric mean of IC_50_ values (left) and the average fold change relative to neutralization of the WA1/2020 virus (right). **(D)** Cross-reactivity of S2X324 with sarbecovirus clades 1a and 1b RBDs analyzed by ELISA. (**E**) Preincubation of serial dilutions of S2X324 or S2E12 with the SARS-CoV-2 RBD prevents binding to the immobilized human ACE2 ectodomain in ELISA. Error bars indicate standard deviation between replicates. (**F**) S2X324-mediated S_1_-shedding from cell surface–expressed SARS-CoV-2 S as determined by flow cytometry. S2E12 and S2X259 were used as positive controls whereas S2M11 and S309 were used as negative controls.

Sotrovimab, a site IV mAb with broad sarbecovirus (clade Ia and Ib) cross-neutralizing activity (*37*), had a moderate (16-fold, 7.3-fold, 21.3-fold, 22.6-fold, 16.6-fold and 8.3-fold) decrease in potency against VSV pseudoviruses expressing BA.2, BA.3, BA.4 (V3G), BA.5, BA.2.12.1 and BA.2.75 S proteins, respectively, compared to BA.1 (2.7-fold), although no additional BA.2, BA.3, BA.4, BA.5 or BA.2.12.1 residue mutations map to the epitope (**Fig. 2A-B**) (*37–39*). Similar IC50 fold shifts were observed against authentic isolates of BA.1, BA.2, BA.5, and BA.2.12.1 (**Fig. 2C** and **Fig. S11**). Importantly, sotrovimab retained in vitro effector functions against BA.2 and conferred Fc-dependent protection in the lungs of mice infected with BA.2 (*40*). The reduced neutralization might result from the S371F substitution, which is found in BA.2, BA.3, BA.4 and BA.5 and introduces a bulky residue that could reposition the nearby N343 glycan, which is part of the sotrovimab epitope, as supported by molecular dynamics simulations (*41*) (**Fig. S9A-E**). Although we could not test the effect of the S371F substitution alone in the Wu-G614 S background (due to poor VSV pseudovirus infectivity), the S371F, S373P and S375F triple mutant (as found in BA.2, BA.3, BA.4, BA.5 and BA.2.12.1) reduced sotrovimab-mediated neutralization by 3.4-fold relative to Wuhan-Hu-1 (D614). Moreover, the S371L, S373P and S375F triple mutant (as found in BA.1) did not alter sotrovimab activity (**Fig. S9F** and **Table S4**), lending further support to the role of F371 in repositioning the N343 glycan and explaining the reduced potency against BA.2 and BA.3. S2X259, a site IIa mAb that broadly reacts with the RBD of multiple sarbecoviruses (*32*) retained activity against BA.1 (*3*). However, S2X259 neutralization was decreased by more than tenfold against BA.2, BA.3, BA.4, BA.5, BA.2.12.1 and BA.2.75 likely due to the detrimental effect of the R408S mutation (*32*). S2H97 is a site V mAb that experienced a 4.7-fold to 10-fold decrease in neutralization potency against Omicron sublineages compared to Wu-G614 despite the absence of mutations present in the epitope or otherwise found to affect binding by DMS, perhaps reflecting differential accessibility to its cryptic epitope in the context of the S trimer (*26*).

## Identification of the pan-variant and ultrapotent neutralizing mAb S2X324

The S2X324 mAb stood out in our panel as its neutralization potency was unaffected by BA.1, BA.2, BA.3, BA.4, BA.5, BA.2.12.1 and BA.2.75 S mutations (**Fig. 2A**). S2X324 bound to BA.1 and BA.2 RBDs with high affinity (**Fig. S10A** and **Table S5**) and cross-reacted with and neutralized all SARS-CoV-2 variants (VSV-pseudoviruses and authentic viruses) tested with IC_50_s below 10 ng/ml **(Fig. 2B-C** and **Fig. S11)**. S2X324 cross-reacted with the sarbecovirus clade 1b Pangolin-GD RBD, but did not recognize more divergent sarbecovirus RBDs **(Fig. 2D)** including SARS-CoV, distinguishing this mAb from previously described broadly neutralizing sarbecovirus mAbs such as S2X259 (*32*), S2K146 (*36*), S2H97 (*26*) and sotrovimab (*37*). Epitope binning experiments showed that S2X324 did not compete with the site I mAb S2K146 or site II mAbs S304 and S2X259 but did compete with the site I mAb S2H14 and the site IV mAb sotrovimab **(Fig. S10B)**. S2X324 inhibited binding of the SARS-CoV-2 RBD to human ACE2 in a concentration-dependent manner, as measured by competition ELISA **(Fig. 2E)** and induced premature shedding of the S_1_ subunit **(Fig. 2F)**. This suggests that blockage of ACE2 binding as well as possible premature S_1_ shedding (*42*) may participate in S2X324-mediated inhibition of SARS-CoV-2.

To evaluate the ability of S2X324 to modulate antibody dependent-phagocytosis or cytotoxicity, we tested whether the mAb could activate Fcγ receptors expressed at the surface of Jurkat cells. Although S2X324 only activated FcγRIIIa but not FcγRIIa in vitro **(Fig. S12 A-B)**, it triggered both antibody-dependent phagocytosis and cytotoxicity following incubation of peripheral blood mononuclear cells and monocytes with SARS-CoV-2 S-expressing cells in presence of S2X324 **(Fig. S12 C-F).**

## Structural basis for S2X324-mediated neutralization

To understand the pan-variant S2X324 inhibitory activity, we determined a cryo-electron microscopy structure of the Omicron BA.1 S ectodomain trimer bound to the S2X324 Fab fragment at 3.1 Å resolution (**Fig. 3A, Fig S13 and Table S6**). In our structure, the Omicron S trimer has two open RBDs and one closed RBD with three bound Fabs. We used focused classification and local refinement of the closed RBD-S2X324 Fab complex to obtain a 3.3 Å structure revealing the molecular details of the binding interface.

**Figure 3:**
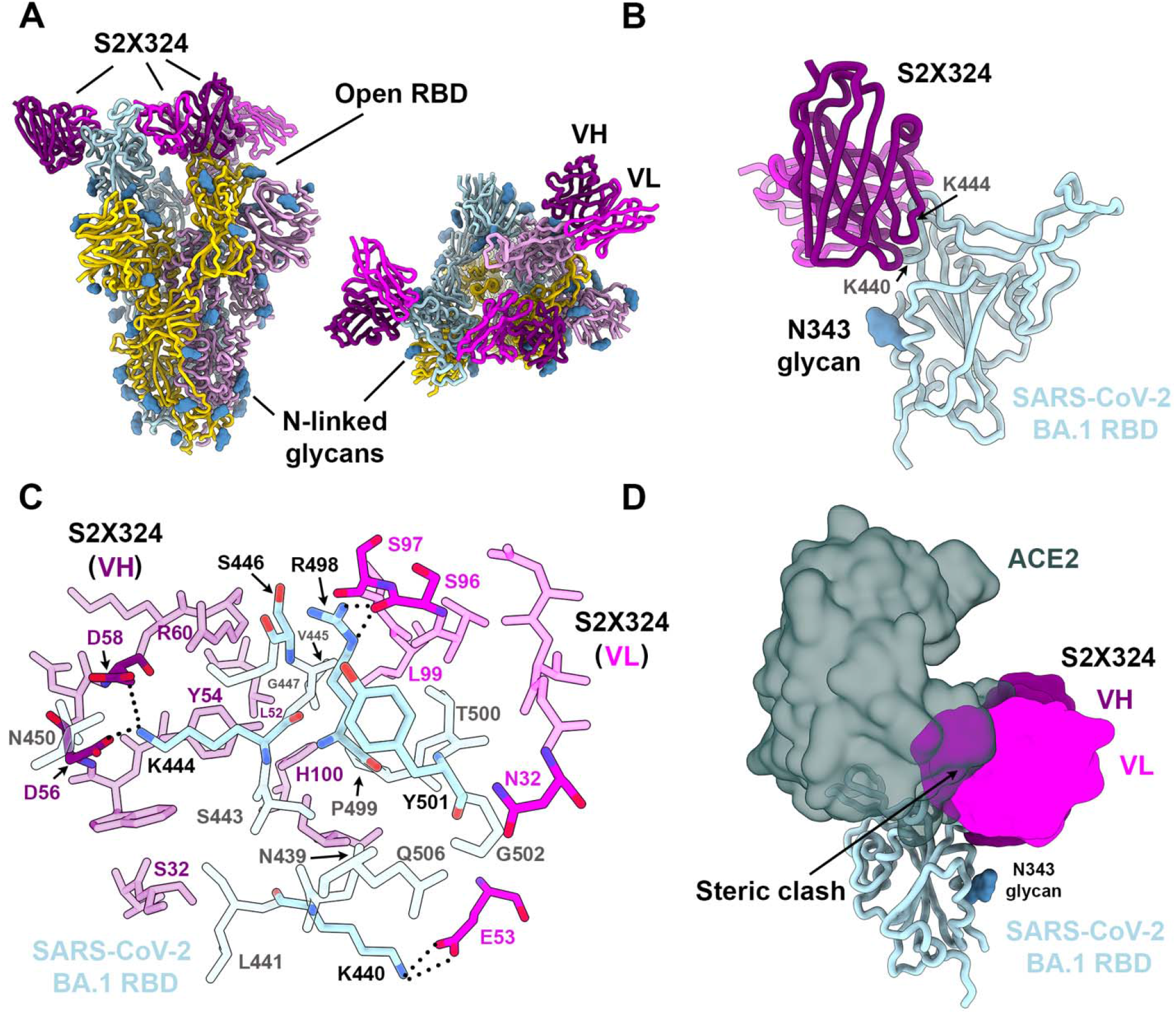
Structural characterization of the S2X324 pan-variant mAb. (**A**) Cryo-EM structure viewed along two orthogonal orientations of the prefusion SARS-CoV-2 Omicron BA.1 S ectodomain trimer with three S2X324 Fab fragments bound. SARS-CoV-2 S protomers are colored light blue, pink, and gold. S2X324 heavy chain and light chain variable domains are colored purple and magenta, respectively. Glycans are rendered as blue spheres. (**B**) Ribbon diagram of the S2X324-bound SARS-CoV-2 RBD. The N343 glycan is rendered as blue spheres. (**C**) Zoomed-in view of the contacts formed between S2X324 and the SARS-CoV-2 RBD. Selected epitope residues are labeled, and electrostatic interactions are indicated with dotted lines. (**D**) Superimposition of the S2X324-bound (purple and magenta) and ACE2-bound [dark gray, PDB 6M0J (*44*)] SARS-CoV-2 RBD (light blue) structures showing steric overlap. The N343 glycan is rendered as blue spheres.

S2X324 recognizes an RBD epitope partially overlapping with antigenic sites Ia/Ib and IV (**Fig. 3A-B)**, explaining the observed competition with S2H14 (*12*) and S309 (*37*) mAbs. S2X324 utilizes all six complementary-determining loops to recognize RBD residues N439, K440, L441, S443, K444, V445, S446, G447, N450, R498, P499, T500, Y501, G502 and Q506 (**Fig. 3C)**. In line with the competition assay (**Fig. 2D)**, S2X324 overlaps with the ACE2-interaction site on the RBD and would sterically hinder receptor engagement (**Fig. 3D)**.

The structure explains how this mAb accommodates residues that are mutated in Omicron lineages relative to Wuhan-Hu-1: N440K (BA.1/BA.2/BA.3/BA.4/BA.5/BA.2.12), G446S (BA.1/BA.3), Q498R (BA.1/BA.2/BA.3/BA.4/BA.5/BA.2.12.1) and N501Y (BA.1/BA.2/BA.3/BA.4/BA.5/BA.2.12.1). Specifically, K440 forms a salt bridge with the VL E53 side chain, S446 forms van der Waals interactions with VH R60 and VL S96/S97 whereas R498 forms electrostatic interactions with the VL S96 backbone. Our structure further suggests that the tighter binding of S2X324 to the Wuhan-Hu-1 and BA.2 RBDs, relative to BA.1 (**Fig. S10A)**, might be due to G446S as although the mutation is clearly accommodated, at least 1 out of 3 favored rotamers for S446 would clash with the Fab. The Y501 backbone forms van der Waals interactions with the VL N32 side chain which are independent of the RBD residue identity at this position (explaining retention of neutralization of all Y501-containing variants). S2X324 and LY-CoV1404 share 87% and 91% amino acid sequence identity in their heavy and light chains, respectively, likely explaining their similar binding mode (*43*) and pan-variant neutralizing activity, including accommodation of mutations at position 452 (as found in BA.4 and BA.5), which are outside the epitope (**Fig. S14**).

## Identification of S2X324 viral escape mutants in vitro

To explore potential mutations that could promote escape from S2X324-mediated neutralization, we passaged a replication-competent VSV chimera harboring either SARS-CoV-2 S Wu-G614 (*45*) or Omicron BA.1 S in the presence of S2X324. Residue substitutions at three distinct sites emerged (**Fig. 3C** and **Fig. S15A-B)**: (i) K444N/T (Wu-G614 and BA.1 background) and K444E/M (BA.1 background), that would abrogate the salt bridges formed between the K444 side chain and the heavy chain D56 and D58 side chains; (ii) V445D (Wu-G614 background) and V445A/F (BA.1 background), which would disrupt Van der Waals contacts with S2X324; and (iii) P499R (Wu-G614 background) and P499S/H (BA.1 background) that might alter the local RBD backbone conformation and/or sterically hinder mAb binding **(Tables S7-S9)**. Furthermore, three additional mutations were retrieved in the BA.1 S background only: S446I, G447S, and N448K, which reside at the interface between the heavy and light chains. The K444N, V445D and P499R S chimeras were outcompeted by the Wuhan-Hu-1/D614G S chimera after four rounds of passaging, suggesting a modest loss of fitness in this replicating chimeric virus model system (**Fig. S15C**). Although each of these mutations require a single nucleotide substitution, they are very rare and as of May 2^th^ 2022 they have been detected cumulatively only in 0.082% and 0.02% of Delta and Omicron genome sequences, respectively **(Table S9** and **Fig. S16A)**. We further tested VSV pseudoviruses (Wu-G614 background) carrying K444E, K444D, K444N and V445D and confirmed that these mutations abrogated S2X324 neutralizing activity **(Fig. S16B** and **Table S10)**. Importantly, S2X324 retained potent neutralizing activity against V445T, G446V and P499S, as well as against other mutations in the epitope found in known variants such as N439K, N440K and N501Y.

## S2X324 protects hamsters against SARS-CoV-2 Delta and BA.2 variants

Using Syrian hamsters challenged with SARS-CoV-2, we investigated the *in vivo* protective efficacy of S2X324 prophylactically and therapeutically. These experiments used S2X324 VH/VL fused to hamster IgG2a constant regions to allow promoting antibody-mediated effector functions in this animal model. Prophylactic administration of S2X324 (blue) or S309 (green) protected hamsters challenged with SARS-CoV-2 Delta in a dose-dependent manner with S2X324 exhibiting an approximately 3-fold higher efficacy than S309 (**Fig. 4A-C**), despite a 20-fold difference observed in terms of in vitro potency against SARS-CoV-2 Delta S VSV(**Fig. 2B**). These data support the lack of direct correlation between in vitro and in vivo potency as previously reported (*40, 46*). Moreover, prophylactic administration of S2X324 hamster IgG2a at 5 mg/kg decreased viral RNA loads in the lungs and trachea of hamsters challenged with BA.2 below detection levels (**Fig. 4D**). Therapeutic S2X324 administration (one day after challenge) at 2 and 5 mg/kg reduced lung viral RNA by 2.5 and 3 orders of magnitude compared to the control group, respectively (**Fig. 4E-F**). Viral replication in the lungs was fully abrogated at 5 and 2 mg/kg of S2X324 and reduced by one order of magnitude for animals treated with 0.5 and 0.1 mg/kg of S2X324. No statistically significant differences were observed for animals receiving an Fc-silenced version of S2X324 (N297A, pink), indicating limited contributions of Fc-mediated effector functions in these experimental conditions.

**Figure 4.**
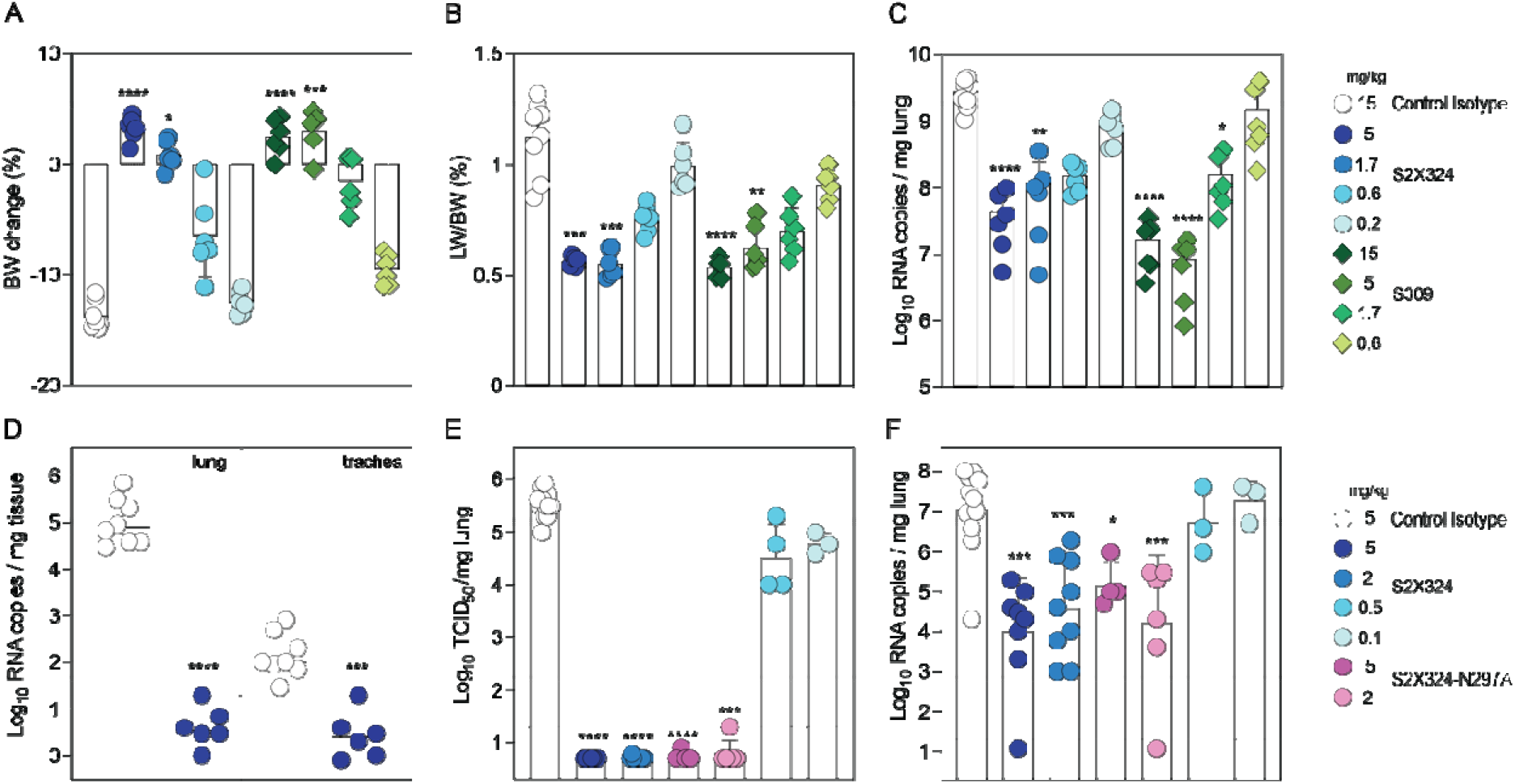
S2X324 protects hamsters against SARS-CoV-2 Delta and BA.2 challenge. **(A-C) A-C,** Dose-dependent (expressed in mg/kg of body weight) prophylactic protection of S2X324 (blue circles) vs S309 (green diamonds) hamster IgG2a (harboring hamster IgG2a constant regions) in animals infected with SARS-CoV-2 Delta evaluated 4 days post infection based on % of body weight change (A), % of lung weight vs body weight ratio (B), viral RNA load (C). (n=6 animals/dose) *, **, ***, **** p< 0.05, p< 0.01, 0.001, and 0.0001 relative to isotype control (MGH2 antibody against circumsporozoite protein of Plasmodium sporozoites), respectively (Mann-Whitney 2-tail T test). **D**, Quantification of viral RNA load in the lung and trachea of Syrian hamsters 4 days after intranasal infection with SARS-CoV-2 Omicron BA.2 which was preceded 1 day prior by prophylactic intraperitoneal administration of S2X324 hamster IgG2a at 5 mg/kg of body weight. **, ***, **** p< 0.01, 0.001, and 0.0001 relative to control, respectively (Kruskal-Wallis test). **E-F**, Quantification of replicating viru titers [50% tissue culture infectious dose (TCID_50_)] (E) and viral RNA load (F) in the lung of Syrian hamsters 4 days after intranasal infection with SARS-CoV-2 Delta followed by therapeutic intraperitoneal administration of S2X324 hamster IgG2a (blue symbols) or N297A mutant IgG2a (purple symbols) at four different doses: 5, 2, 0.5 or 0.1 mg/kg of body weight. *, **, ***, **** p< 0.05, p< 0.01, 0.001, and 0.0001 relative to control, respectively (Mann-Whitney 2-tail T test).

## Discussion

Immunological imprinting, which is also defined as original antigenic sin, was described based on the observation that infections with influenza virus strains similar to that of a prior infection preferentially boost antibody responses against epitopes shared with the original strain (*47*). Although this phenomenon is often considered as detrimental, it can also be beneficial, as was the case after the 2009 H1N1 pandemic: initial antibody responses to infection with this newly emerged and antigenically shifted virus were dominated by antibodies targeting the conserved hemagglutinin stem region (*48, 49*) whereas secondary exposures over the years through vaccination or infection elicited antibody responses to the shifted variant (i.e., to “non-conserved” hemagglutinin epitopes) (*48, 50*). Moreover, several studies reported stem-directed antibody-mediated protection against H5N1 and H7N9 zoonotic influenza strains through hemagglutinin imprinting during childhood resulting from exposure to seasonal H1N1 and H3N2, respectively (*49, 51*). Similarly, we show that exposure to the antigenically shifted Omicron primarily leads to a recall of existing memory B cells specific for epitopes shared by multiple SARS-CoV-2 variants rather than by priming naïve B cells recognizing Omicron-specific epitopes (at least up to 100 days post breakthrough infection), as also recently reported (*52*). Although immune imprinting may be beneficial in stimulating responses to cross-reactive SARS-CoV-2 epitopes, antibody responses to some Omicron-specific epitopes were markedly diminished by prior antigenic exposure.

Currently, there is uncertainty regarding the need for vaccines matching dominant circulating SARS-CoV-2 variants (like those used for seasonal influenza) or if the repeated use of Wuhan-Hu-1-based vaccines will suffice. Recent work showed that boosting previously immunized macaques with Beta or Omicron mRNA S vaccines or with Beta RBD nanoparticle vaccines elicited comparably high titers of antibodies broadly neutralizing multiple variants relative to Wuhan-Hu-1-based vaccines (*53–55*). Furthermore, administration of Wuhan-Hu-1-based vaccine boosters in humans was shown to elicit appreciable titers of neutralizing antibodies and prevent severe disease associated with Omicron infections (*10, 18, 21, 56–58*). The limited cross-variant neutralization elicited by Omicron primary infection in humans or vaccination of immunologically naïve animals and the data on the specificity of memory B cells presented here indicate that an Omicron-based vaccine might only elicit narrow antibody responses directed towards the vaccine-matched antigen, suggesting that a heterologous prime-boost or a multivalent approach might be preferable (*53, 59–66*). Omicron infection (and presumably Omicron S-based vaccination) of previously immune subjects, however, recalls cross-reactive memory B cells which may further mature overtime to enhance their affinity and neutralizing potency against Omicron, but also to broaden their neutralizing activity against past and future variants. Indeed, multiple studies showed that somatic hypermutations yield RBD-specific mAbs with increased affinity for the homotypic antigen and with augmented resilience to immune evasion of emerging variants (*36, 67–70*).

Understanding antibody responses elicited by and directed towards Omicron sublineages is key to inform public health policies and the design of SARS-CoV-2 and sarbecovirus vaccines (*64, 65, 71–73*). Our data show that Omicron breakthrough infections did not elicit high titers of pan-sarbecovirus neutralizing antibodies (e.g., directed against SARS-CoV), in agreement with recent data (*74*). These findings contrast with the observation that pre-existing immunity to SARS-CoV followed by SARS-CoV-2 vaccination was associated with elicitation of pan-sarbecovirus neutralizing antibodies (*27*). These different outcomes might be explained by the low frequency of memory B cells encoding for neutralizing antibodies targeting antigenic sites shared between pre-Omicron variants (Wuhan-Hu-1-related strains), Omicron and SARS-CoV, due to the genetic distance between these three antigenically distinct viruses. For instance, Omicron BA.1 and BA.2 harbor variations of the RBD antigenic site II, which is the target of pan-sarbecovirus neutralizing antibodies such as S2X259 (*32*) and ADG2 (*75*), that lead to resistance to neutralization mediated by some of the mAbs recognizing this site (*3, 17*). This suggests that conservation of RBD sites across sarbecoviruses may have resulted (at least partially) from limited immune pressure rather than from functional or structural constraints (i.e., some mutations at these conserved sites may remain compatible with viral fitness) (*74*).

Finally, recent preclinical assessment of intranasally administered influenza and sarbecovirus vaccine candidates demonstrated the induction of lung-resident protective mucosal humoral and cellular immunity at the site of viral entry (*76–79*). These observations, along with our findings that SARS-CoV-2 breakthrough infections, but not vaccination-only, elicited neutralizing activity in the nasal cavity motivate the development and evaluation of a next generation of vaccines administered intranasally.

## Acknowledgements

This study was supported by the National Institute of Allergy and Infectious Diseases (DP1AI158186 and HHSN272201700059C to D.V.), a Pew Biomedical Scholars Award (D.V.), an Investigators in the Pathogenesis of Infectious Disease Awards from the Burroughs Wellcome Fund (D.V.), Fast Grants (D.V.), the University of Washington Arnold and Mabel Beckman cryoEM center and the National Institute of Health grant S10OD032290 (to D.V.). D.V. is an Investigator of the Howard Hughes Medical Institute. O.G. is funded by the Swiss Kidney Foundation. For the purpose of open access, the author has applied a CC BY public copyright license to any Author Accepted Manuscript version arising from this submission. We thank Abigail E. Powell and Nadine Czudnochowski for assistance with protein production.

## Author contributions

A.C.W., A.L., D.P., D.C., M.S.P. and D.V. designed the experiments; A.C.W., A.D.M., D.P., C.S., W.R., K.S.S. F.Z., H.V.D., M.G., G.Sc. and F.A.L. isolated mAb and performed binding, neutralization assays, biolayer interferometry and surface plasmon resonance binding measurements; A.R., J.Z., N.F., M.M.R., J.N. performed neutralization assays using authentic virus. H.K. confirmed the Spike mutations of authentic virus by Sanger sequencing. A.D.M. and D.P. performed ACE2 binding inhibition and S_1_ shedding assays; B.G. and M.A.S. evaluated effector functions; C.S.F., J.B. and L.P performed memory B cell repertoire analysis. O.G., A.C. and P.F. contributed to the recruitment of donors and collection of plasma samples. J.d.I. and L.S. performed bioinformatic and epidemiology analyses. Z.L. and S.P.J.W. performed mutant selection and fitness assays; R.A., J.J., F.B., P.M., J.N., G.D.d.M., L.K. and H.B. performed hamster model experiments and data analysis; Y.J.P. carried out cryoEM specimen preparation, data collection and processing. Y.J.P. and D.V. built and refined the atomic models. J.E.B., C.S. purified recombinant glycoproteins. A.L., D.P., Y.J.P., A.D.M., Z.L., F.A.L., D.P., D.C., M.S.P. and D.V. analyzed the data; A.C.W., D.P., D.C., M.S.P. and D.V. wrote the manuscript with input from all authors; F.B., G.S., J.N., S.P.J.W., H.W.V, M.S.P., D.C., and D.V. supervised the project.

## Data and Materials Availability

The cryoEM map and coordinates have been deposited to the Electron Microscopy Databank and Protein Data Bank with accession numbers TBD. Materials generated in this study will be made available on request, but may require a completed materials transfer agreement signed with Vir Biotechnology Inc. or the University of Washington.

---

**Fig. S1A:**
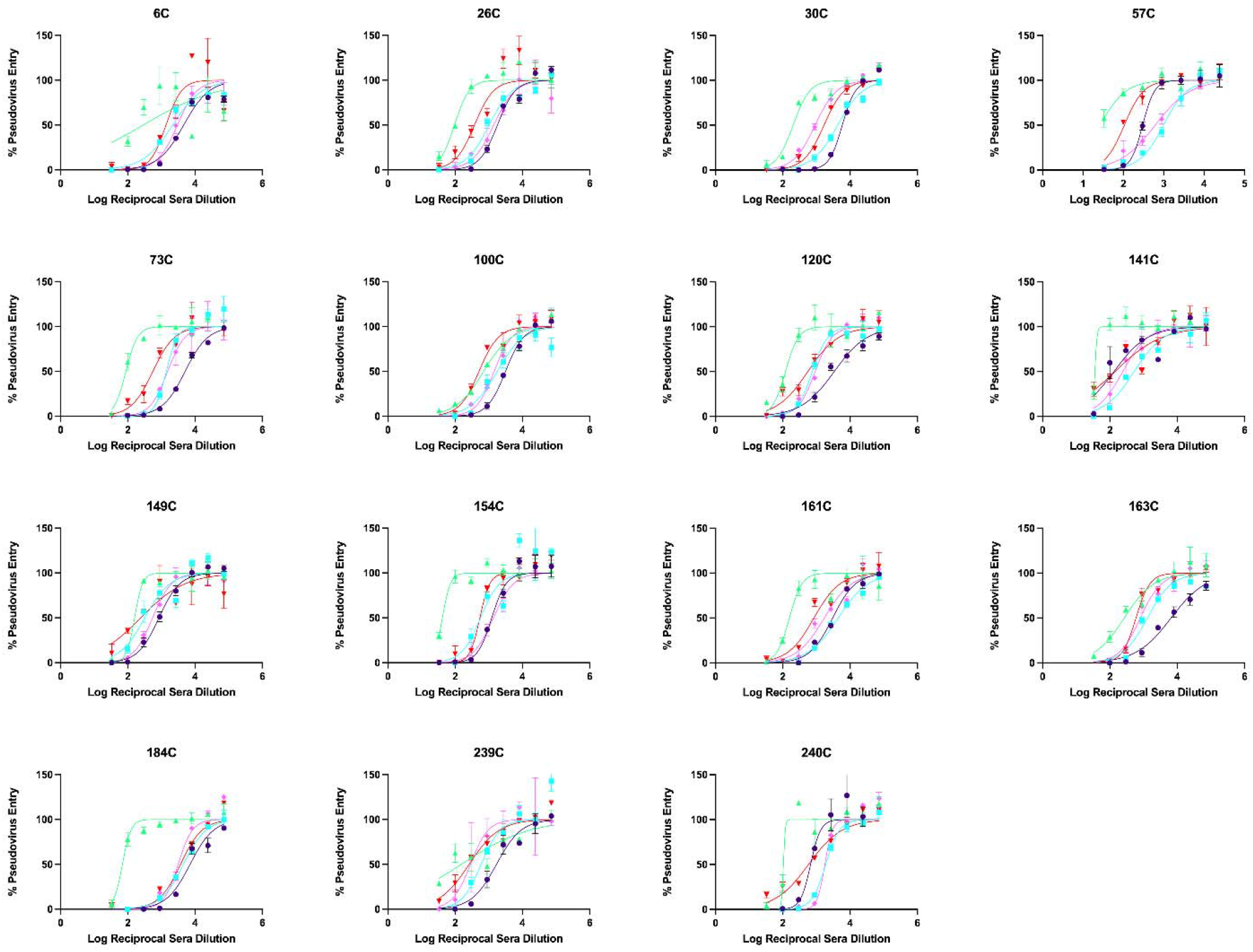
Infected-vaccinated plasma VSV neutralization curves. Purple shows G614, Teal is Delta, Red is Omicron BA.1, Light Pink is Omicron BA.2, and Green is SARS-CoV.

**Fig. S1B:**
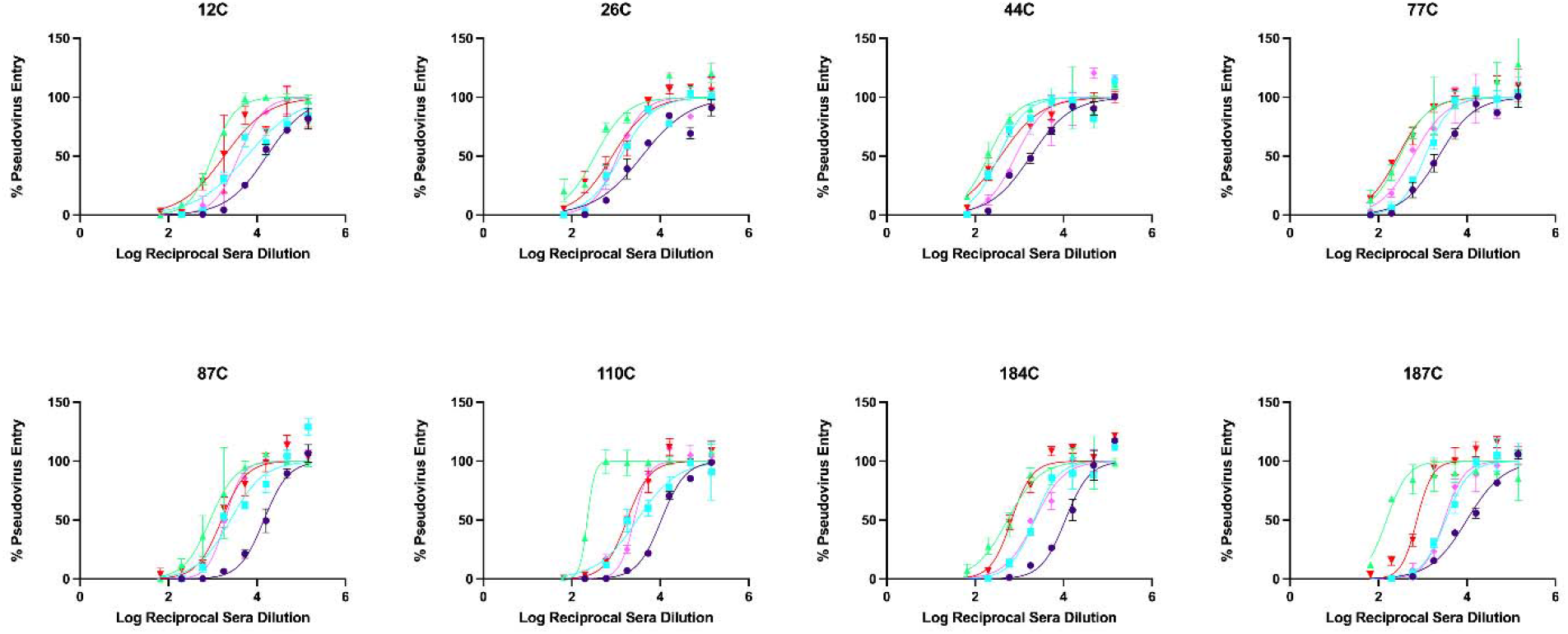
Delta breakthrough plasma VSV neutralization curves. Purple shows G614, Teal is Delta, Red is Omicron BA.1, Light Pink is Omicron BA.2, and Green is SARS-CoV.

**Fig. S1C:**
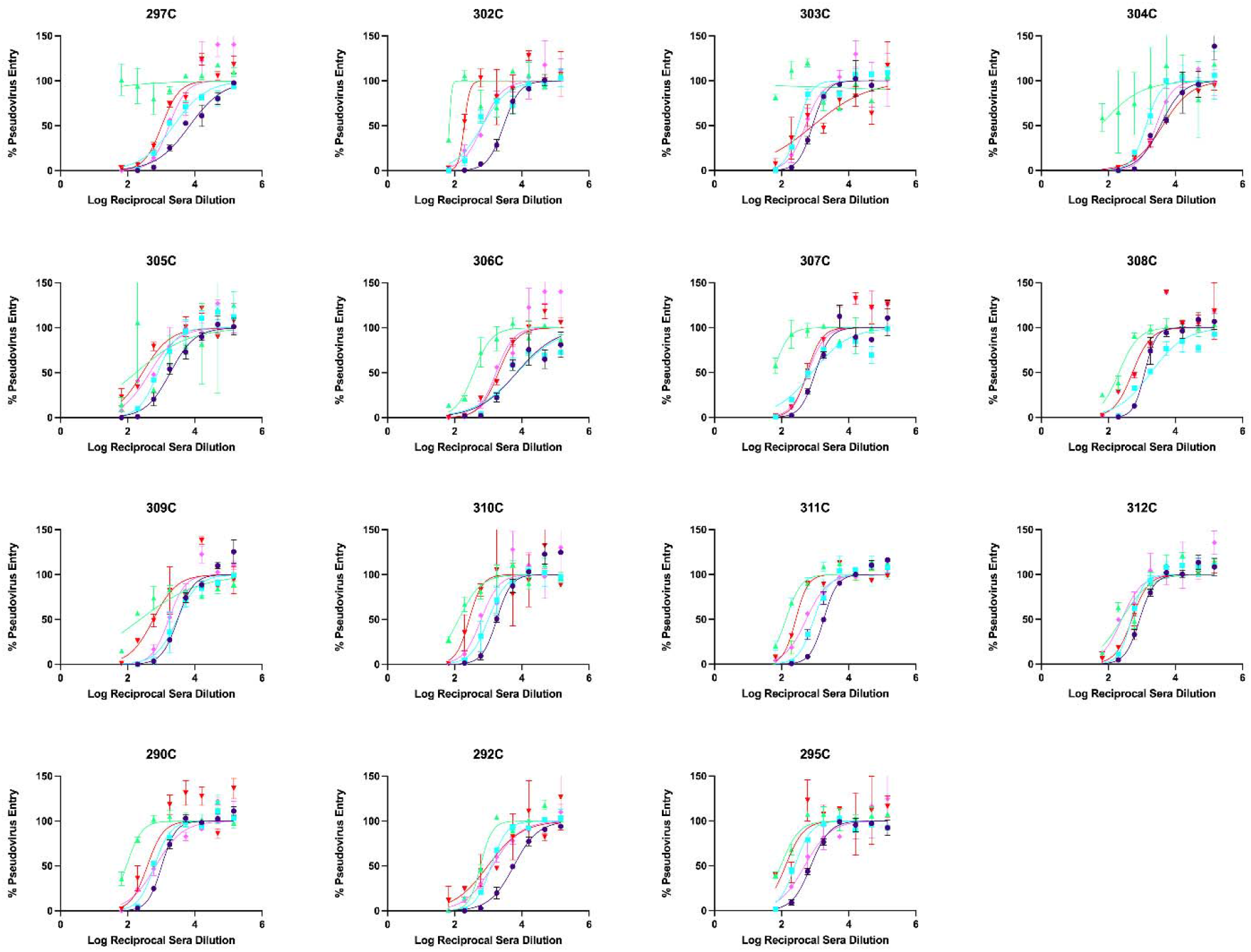
Omicron BA.1 breakthrough plasma VSV neutralization curves. Purple shows G614, Teal is Delta, Red is Omicron BA.1, Light Pink is Omicron BA.2, and Green is SARS-CoV.

**Fig. S1D:**
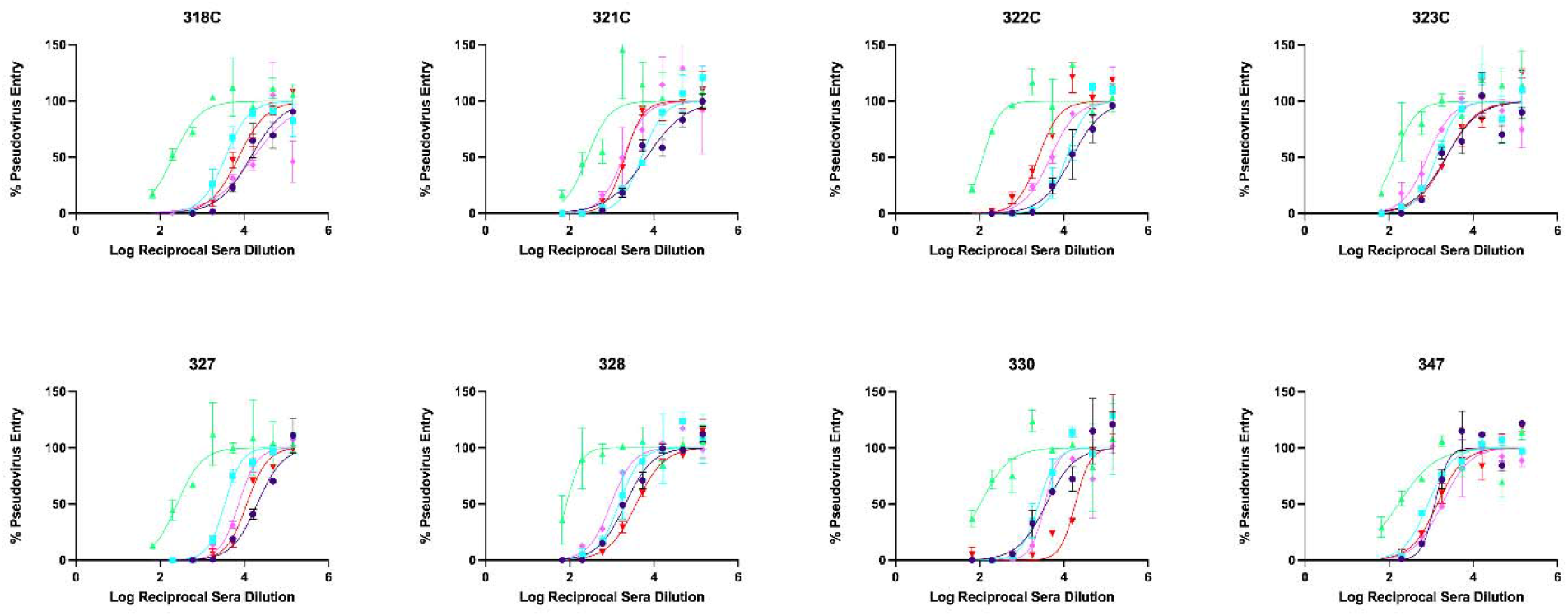
Infected-boosted plasma VSV neutralization curves. Purple shows G614, Teal is Delta, Red is Omicron BA.1, Light Pink is Omicron BA.2, and Green is SARS-CoV.

**Fig. S1E:**
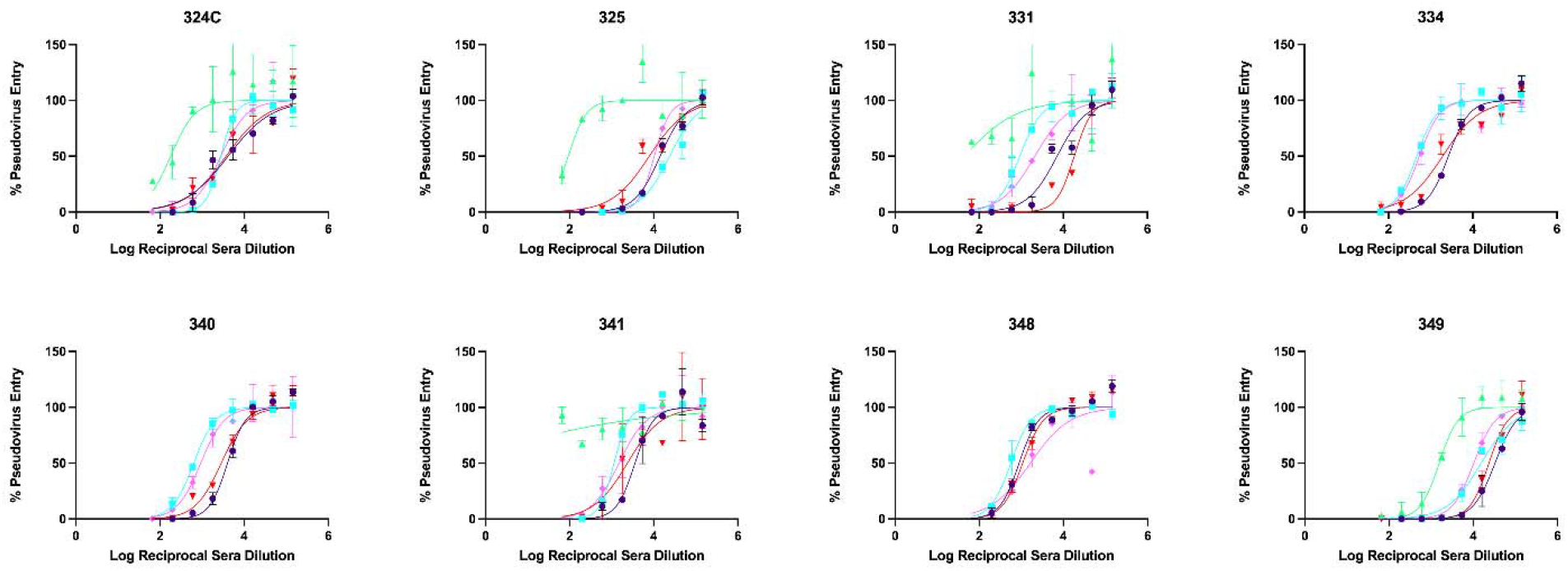
Vaccinated only-boosted plasma VSV neutralization curves. Purple shows G614, Teal is Delta, Red is Omicron BA.1, Light Pink is Omicron BA.2, and Green is SARS-CoV.

**Fig. S1F:**
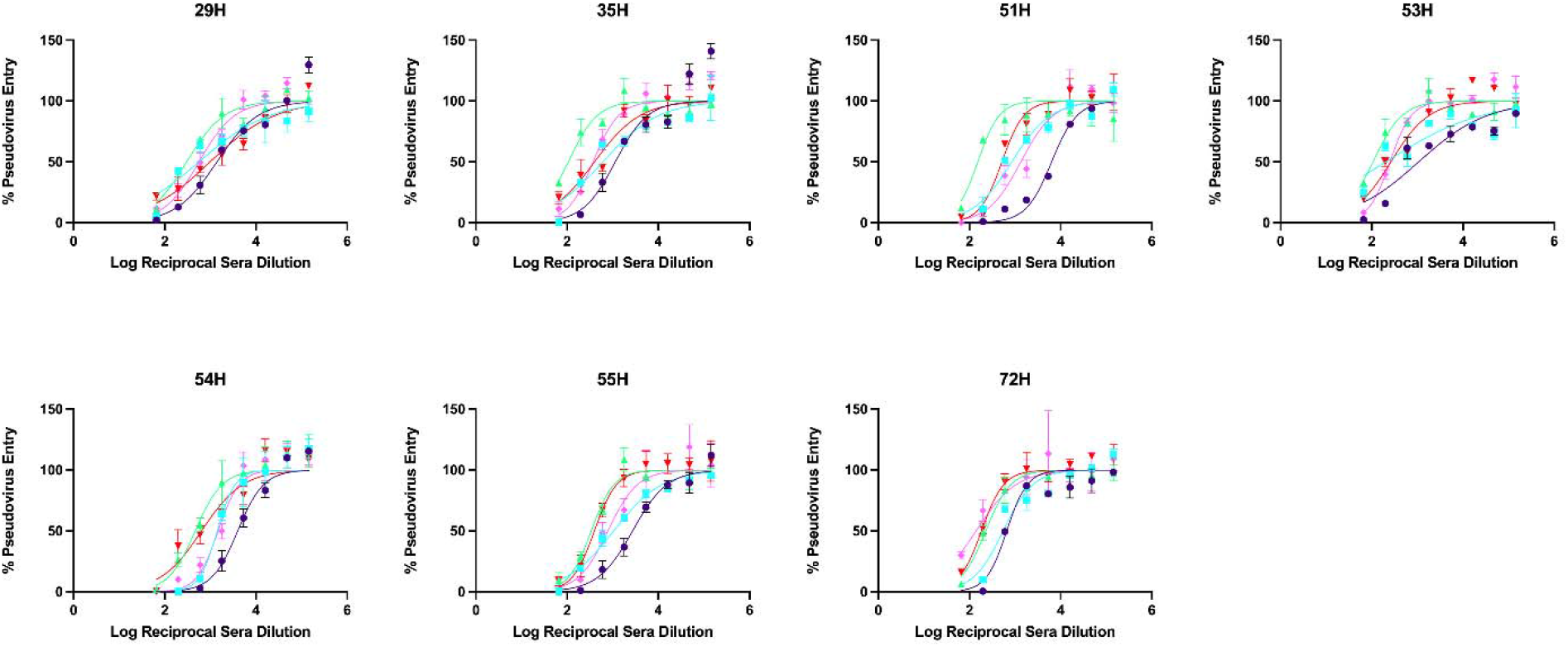
Omicron BA.1 breakthrough boosted plasma VSV neutralization curves. Purple show G614, Teal is Delta, Red is Omicron BA.1, Light Pink is Omicron BA.2, and Green is SARS-CoV.

**Fig. S2.**
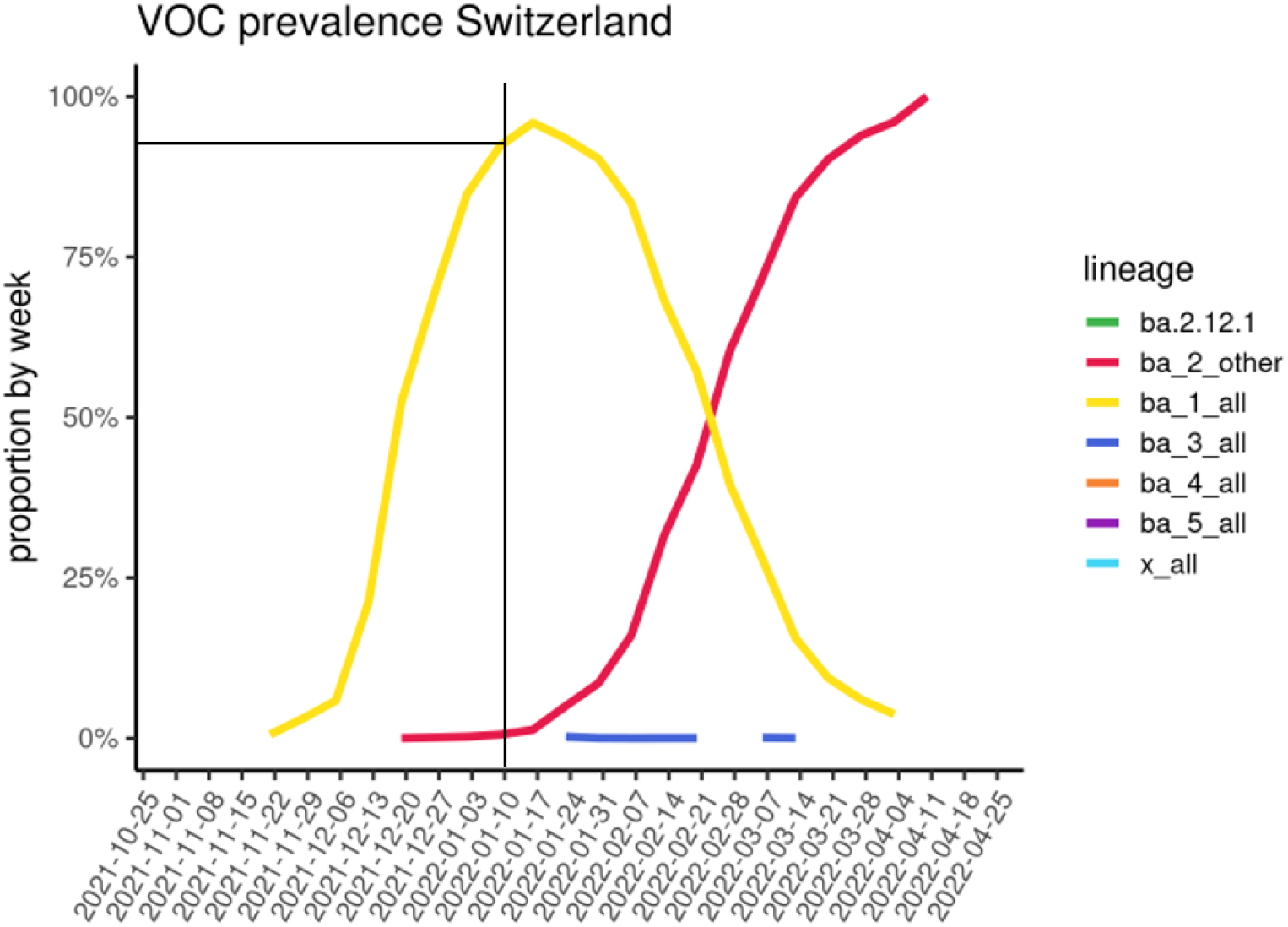
Variant prevalence in Switzerland at the end of 2021 and early 2022. **Data S1.** Description of the Cohorts of SARS-CoV-2-immune, vaccinated, or hybrid individuals, Related to Figure 1A, E-F (External PDF)

**Fig. S3.**
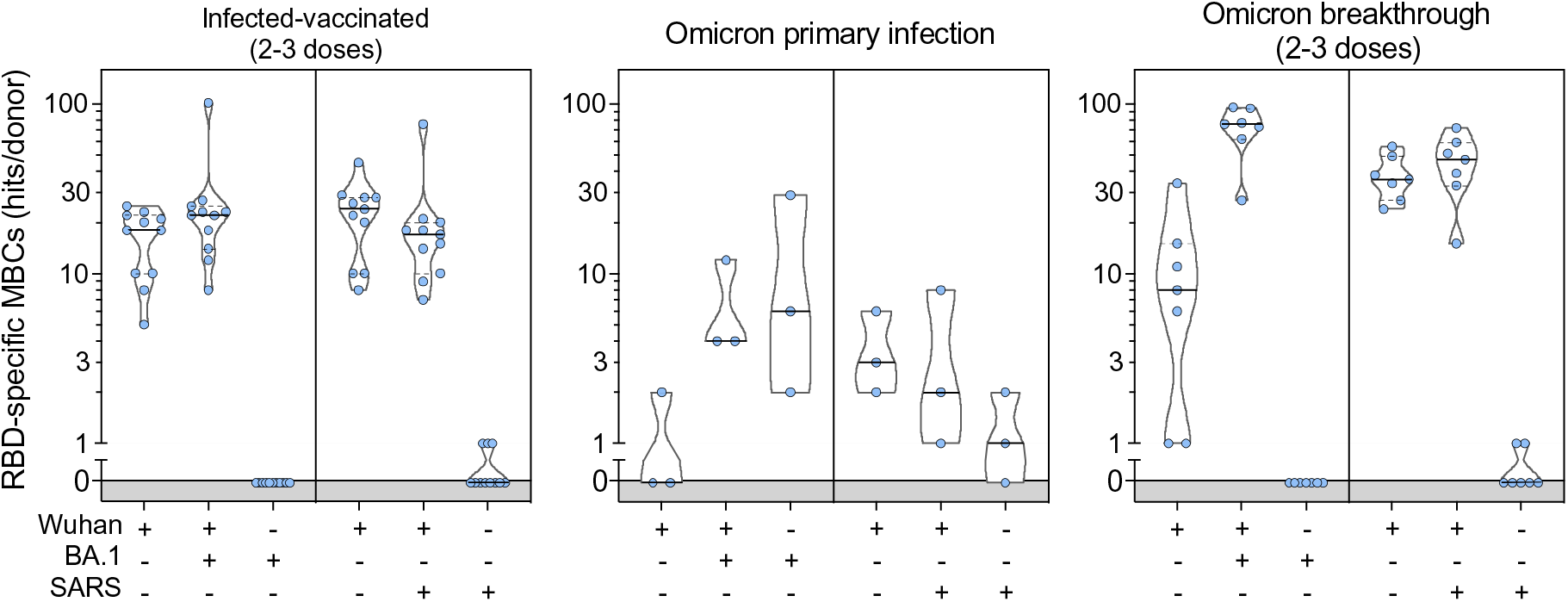
Cross-reactivity of IgGs secreted from memory B cells obtained from infected-vaccinated individuals (left, n=11), subjects who experienced a primary infection with Omicron (center, n=3) or a breakthrough infection in January-March 2022 (right, n=7). Individual dots correspond to number of positive hits as shown in Fig. 1B relative to individual donors.

**Fig. S4.**
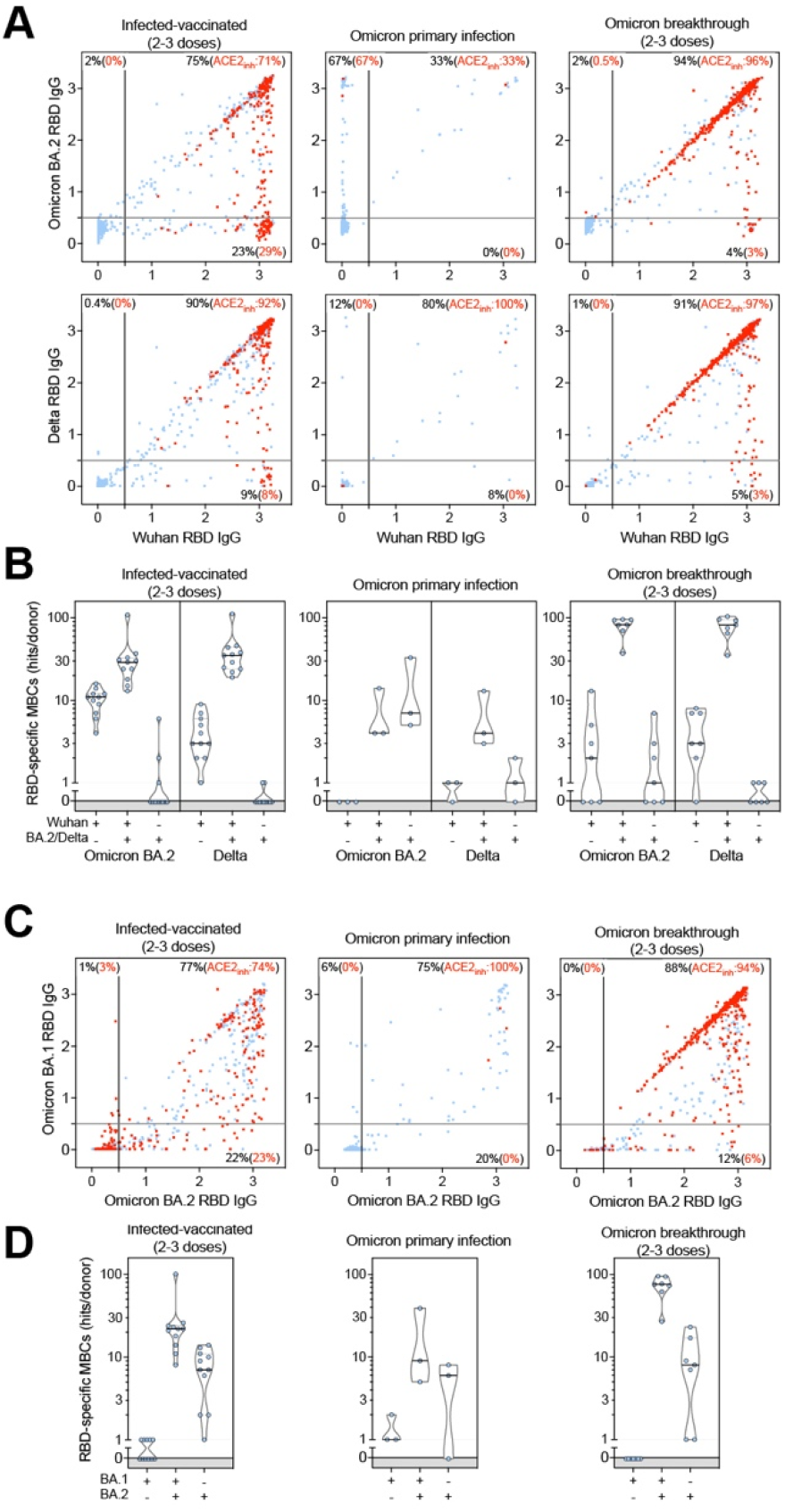
Cross-reactivity of IgG antibodies derived from memory B cells. Antigen-specific memory B cell repertoire analysis (AMBRA) on PBMC obtained from infected-vaccinated individuals (left, n=11), primary SARS-CoV-2 infection (center, n=3) or breakthrough cases (right, n=7) occurring in January-March 2022 (Table S2). (**A** and **C)**, Each dot represents a well containing oligoclonal B cell supernatant screened for the presence of secreted IgGs binding to SARS-CoV-2 Wuhan-Hu-1 and BA.2 RBDs or Wuhan-Hu-1 and Delta RBDs using ELISA (A). A similar analysis on the same samples for the presence of secreted IgGs binding to the SARS-CoV-2 BA.1 and BA.2 RBDs (C). Red dots indicate inhibition of the interaction with ACE2 (using Wuhan-Hu-1 target antigen) as determined in a separate assay (Fig. S5). Percentages are relative to the total of positive hits against any of the antigen tested. **(B** and **D)**, Number of positive hits in panels (A, C) relative to individual donors.

**Fig. S5.**
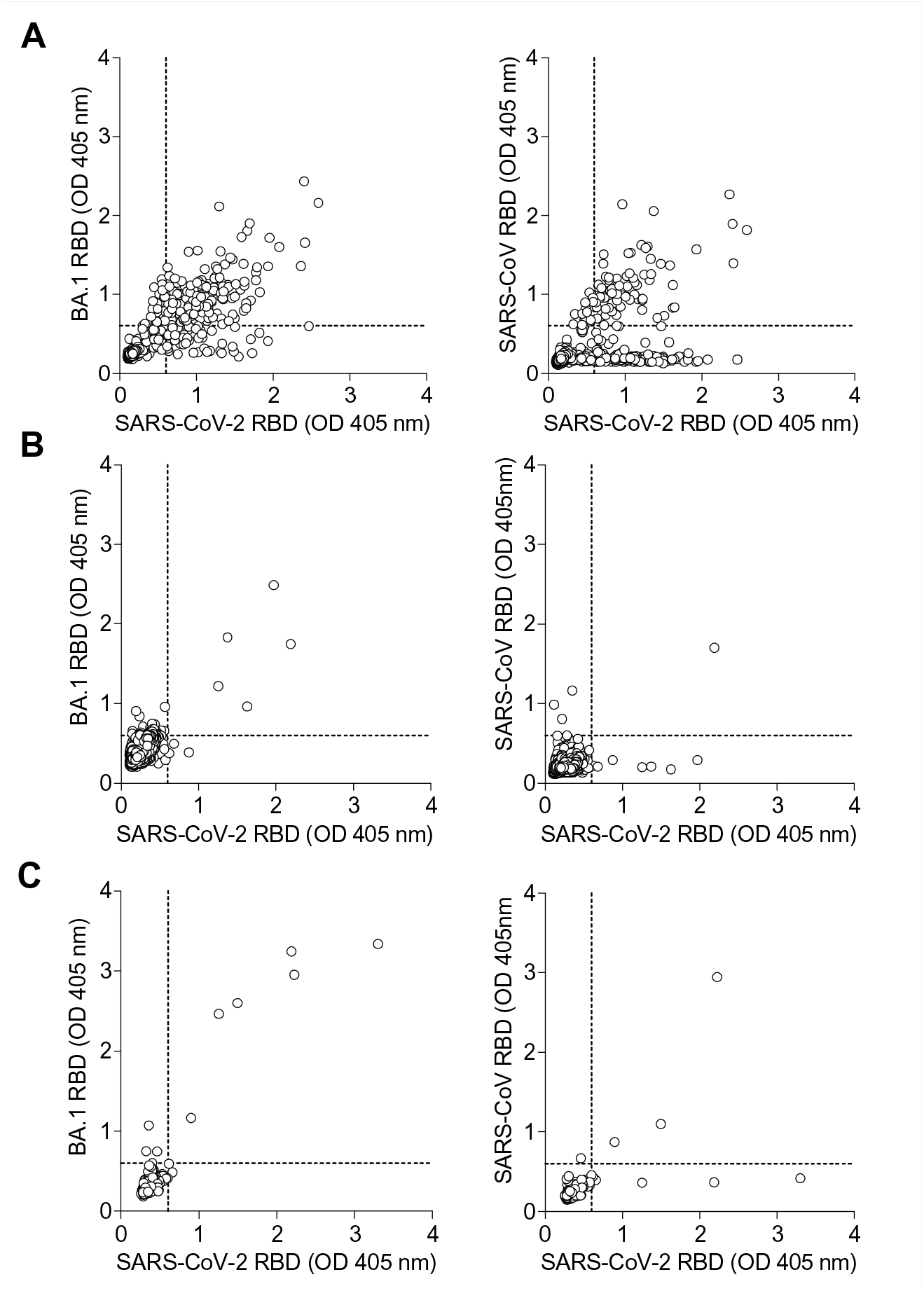
Cross-reactivity of antibodies obtained from single-plasma cell cultures. Circulating CD138+ cells isolated from 3 vaccinated donors (A to C) approximately 1 week after Omicron breakthrough infection and cultured at 0.5 cells/well for 3 days. Culture supernatants were tested by ELISA for the presence of IgG binding to the SARS-CoV-2 Wuhan-Hu-1, BA.1 and SARS-CoV RBDs. Each dot represents the optical density (OD) values of individual cultures for each antigen tested.

**Fig. S6.**
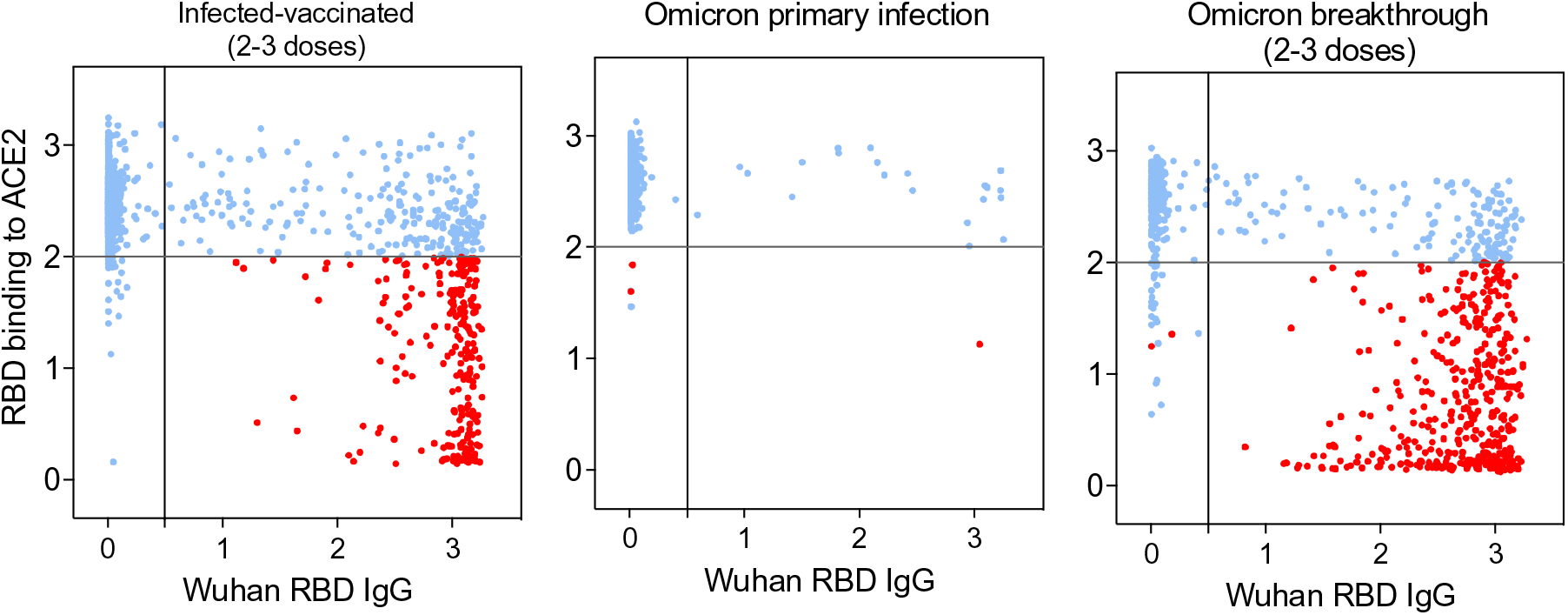
Inhibition of RBD binding to human ACE2 by memory B cell-derived secreted antibodies. Antibody-mediated inhibition of SARS-CoV-2 Wuhan-Hu-1 RBD binding to solid phase ACE2 in function of IgG binding to matched RBD, as determined by ELISA. Each dot represents the optical density (OD) values of individual cultures. Red: cultures with antibodies inhibiting ACE2 binding to the RBD (as defined by a y-axis cutoff of OD≤2)

**Fig. S7.**
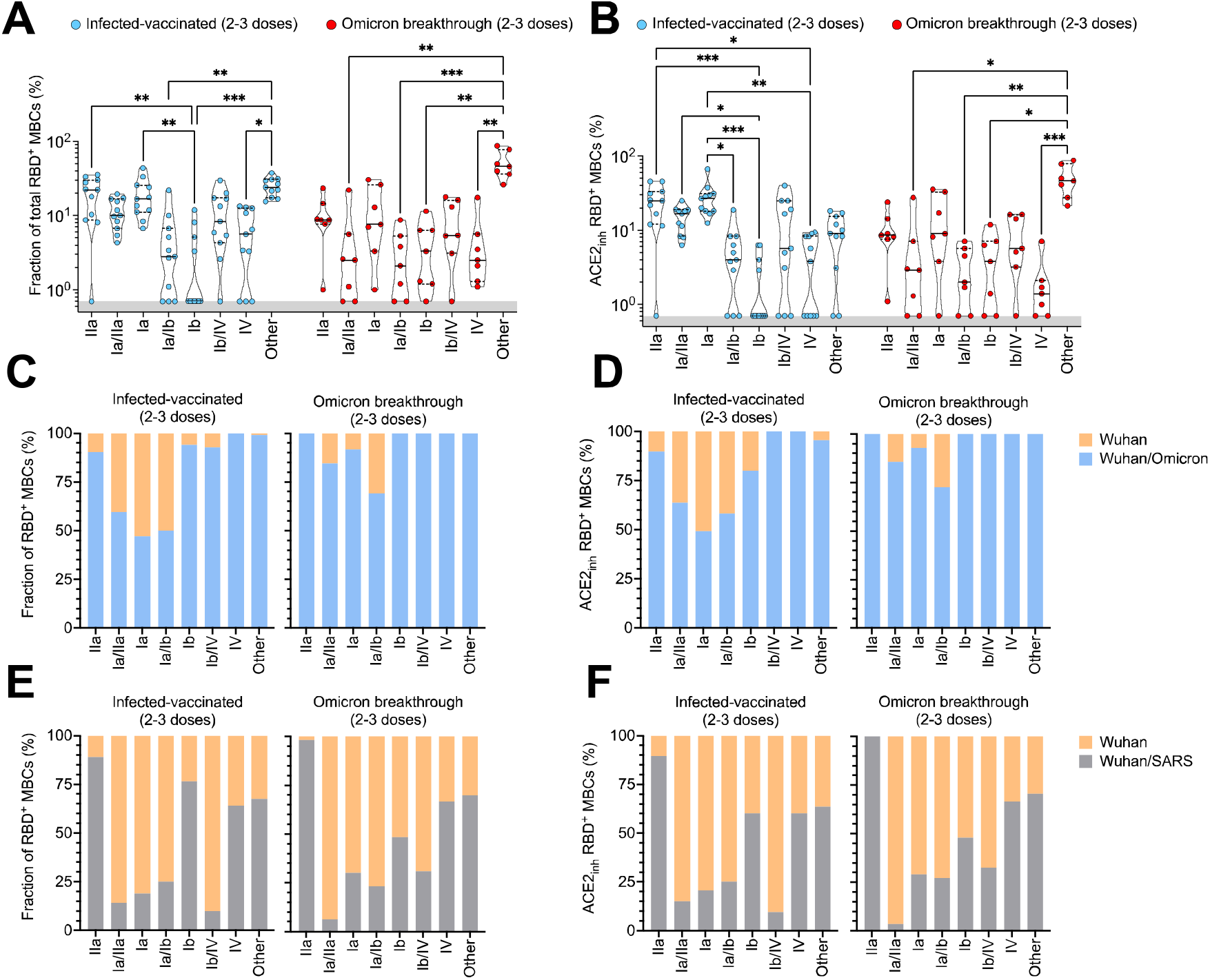
RBD site definition and frequency analysis of IgG antibodies derived from memory B cells. RBD sites targeted by IgG derived from memory B cells were defined by a blockade-of-binding assay using monoclonal antibodies specific for sites Ia (S2E12), Ib (S2X324), IIa (S2X259) and IV (S309). Hybrid sites Ia/IIa, Ia/Ib and Ib/V were also defined when block of binding was observed by both corresponding mAbs. Lack of blockade by probe mAbs is indicated as “Other”. **A-F** Graphs show frequency of total RBD-specific IgGs (A, C, E) and those inhibiting binding of RBD to human. Violin plots (A, B) show percentages of site-specific IgGs in individual donors analyzed. Bars (C-F) show cumulative frequencies of site-specific IgG antibodies filtered for their binding to Wuhan-Hu-1, Omicron BA.1/BA.2 and SARS-CoV RBDs.

**Fig. S8.**
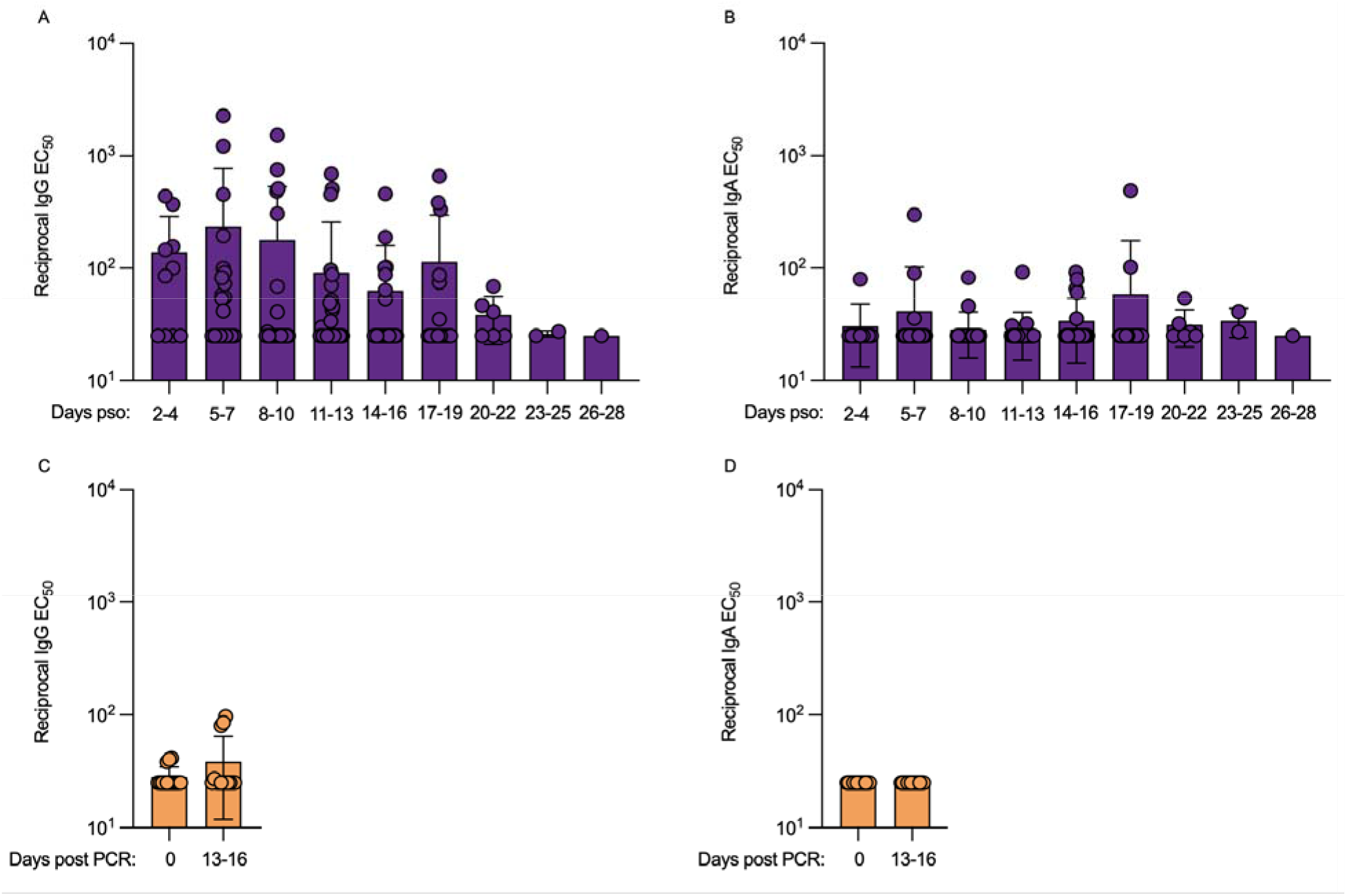
IgG (A, C) or IgA (B,D) binding titers evaluated by ELISA (antigen: Wu-G614 S V-FLIP) in nasal swabs obtained longitudinally upon BA.1 breakthrough infection post symptom onset (pso, A,B) or in vaccinated-only individuals following a negative PCR test.

**Fig. S9.**
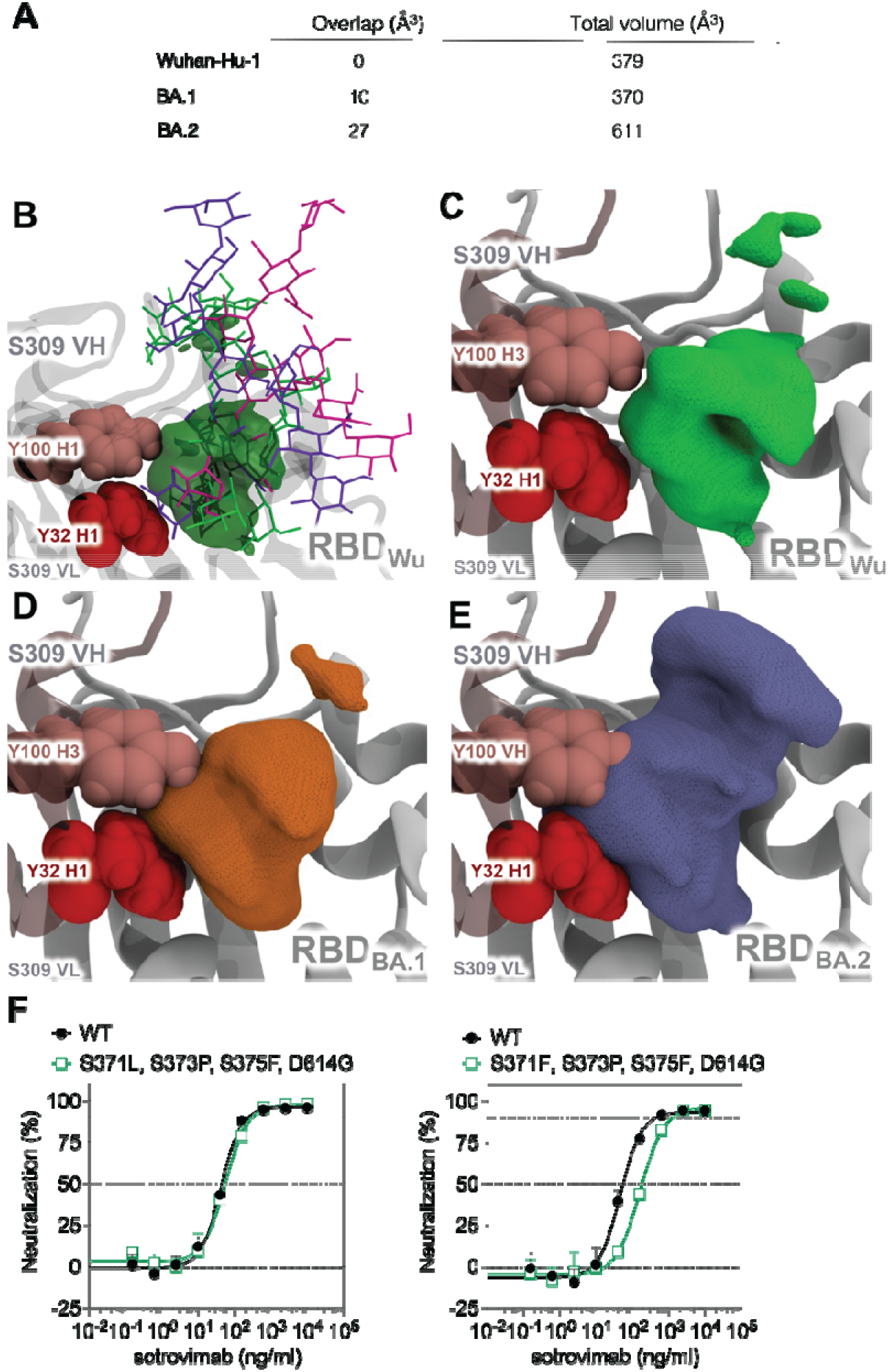
BA.1 and BA.2 mutations impact the conformational preference of RBD glycans in MD simulations. (**A**) Summary of the overlap between the glycan and S309 volumetric density maps (Overlap) along with the total volume occupied by the glycan in the volumetric density map (Total volume). Volumetric density maps quantify the space occupied by the glycan with a maximum probability (isovalue) oS§f 0.057 (see Methods) during an MD simulation; smaller isovalues increase the volume size because ephemeral glycan conformations are included in the map whereas a larger isovalue reduces the volume size because only persistent glycan conformations are included in the map. (**B**) Overlay of three representative glycan conformations from MD of the RBD (Wuhan-Hu-1) shown as sticks, the volumetric map in translucent green, Y32 in H1 and Y100 in H3 as red spheres. (**C**) Volumetric density map of the glycan in RBD (Wuhan-Hu-1) is shown in green, the VH and VL of S309 is shown as a translucent cartoon, Y32 and Y100 in VH as spheres, and the RBD a grey cartoon. (**D**) Volumetric density map of the glycan in RBD (BA.1) is shown in orange; otherwise depicted as in (**C**). (**E**) Volumetric density map of the glycan in RBD (BA.2) is shown in blue; otherwise depicted as in (**C**). Volumetric density maps are asymmetric because glycan dynamics are heterogeneous. (**F**) Neutralizing activity of sotrovimab against VSV pseudoviruses harboring the SARS-CoV-2 Wuhan-Hu-1 S (D614) with the indicated substitutions.

**Fig. S10.**
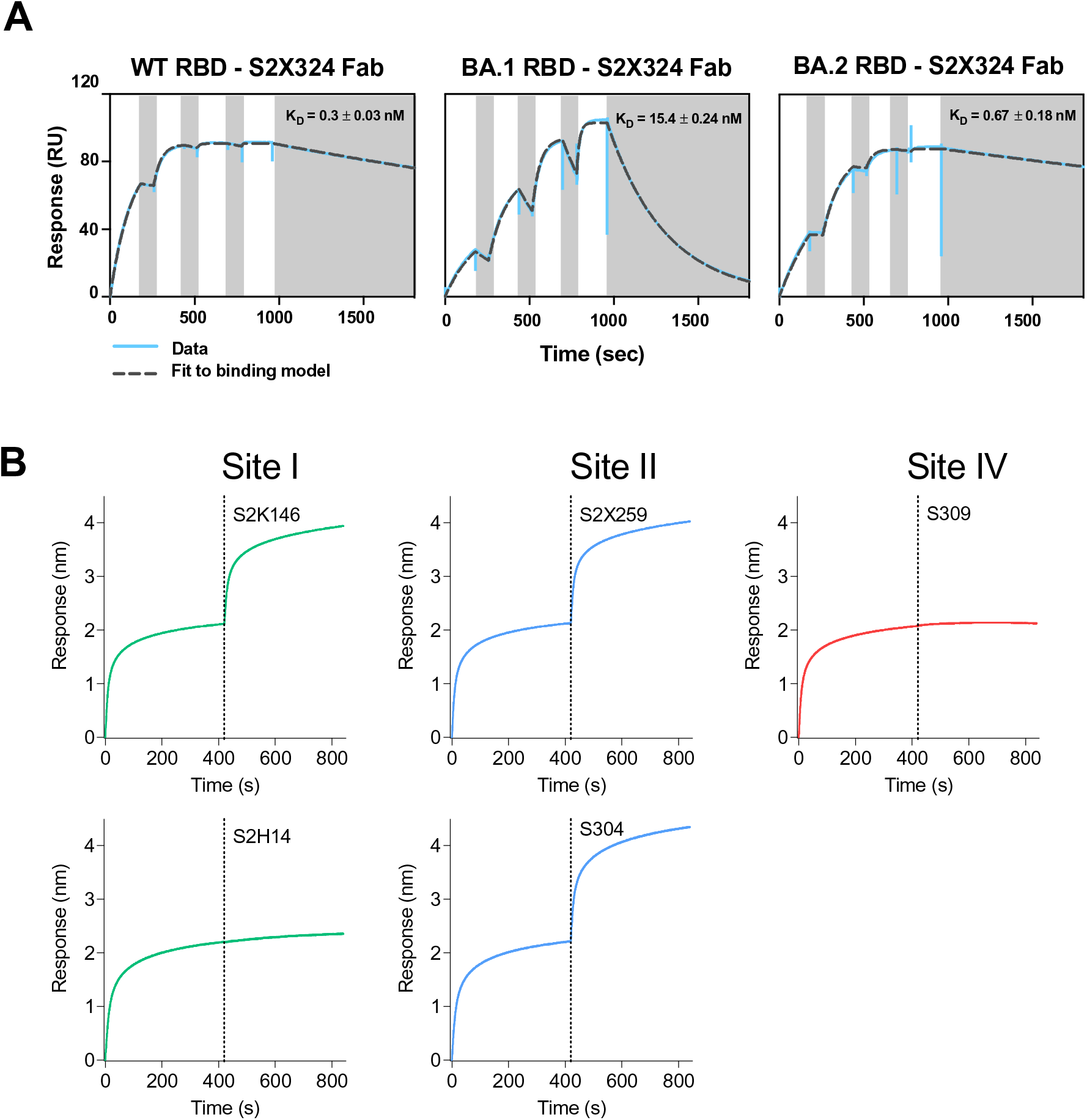
Binding affinity and site specificity of S2X324 mAb. (**A**) Binding of the S2X324 Fab to the SARS-CoV-2 Wuhan-Hu-1 (Wild-type, WT) RBD, Omicron (BA.1) RBD or Omicron BA.2 RBD immobilized at the surface of SPR chips. Experiments were performed with a 3-fold dilution series of Fab: 300, 100, 33, 11 nM and were run as single-cycle kinetics. (**B**) Competition binding assays for S2X324 versus site-I-targeting S2K146, S2H14; site-II targeting S2X259, S304; and site-IV-targeting S309 mAbs on SARS-CoV-2 RBD as measured by biolayer interferometry. One independent experiment out of two is shown.

**Fig. S11.**
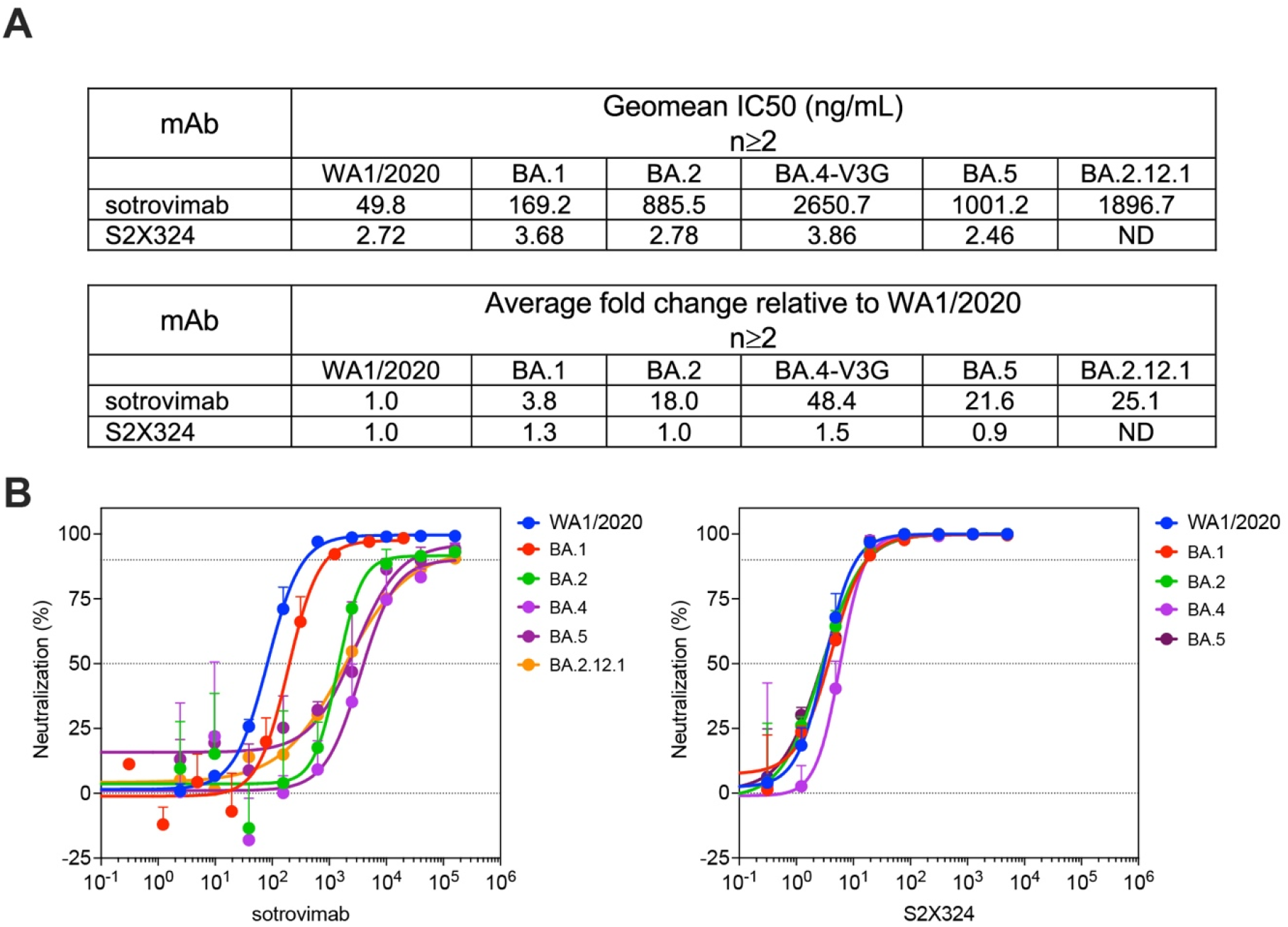
Neutralization of authentic SARS-CoV-2 variants by sotrovimab and S2X324. (**A**) Geometric mean of IC50 values and average fold change for the neutralization of Omicron sublineages and WA1/2020 by sotrovimab and S2X324 mAbs. Strains tested: USA-WA1/2020; BA.1: hCoV-19/USA/MD-HP20874/2021; BA.2: hCoV-19/USA/MD-HP24556/2022: BA.4: hCoV-19/USA/MD-HP30386/2022; BA.5: hCoV-19/USA/COR-22-063113/2022; BA.2.12.1: USA/NY-MSHSPSPPV56475/2022 (B) Representative curves showing neutralization of SARS-CoV-2 strains by sotrovimab and S2X324. Data represent the means of triplicates ± standard deviation from one experiment. Omicron BA.1 data are reported from (*39*). Graph shown is representative of at least 2 independent experiments.

**Fig. S12.**
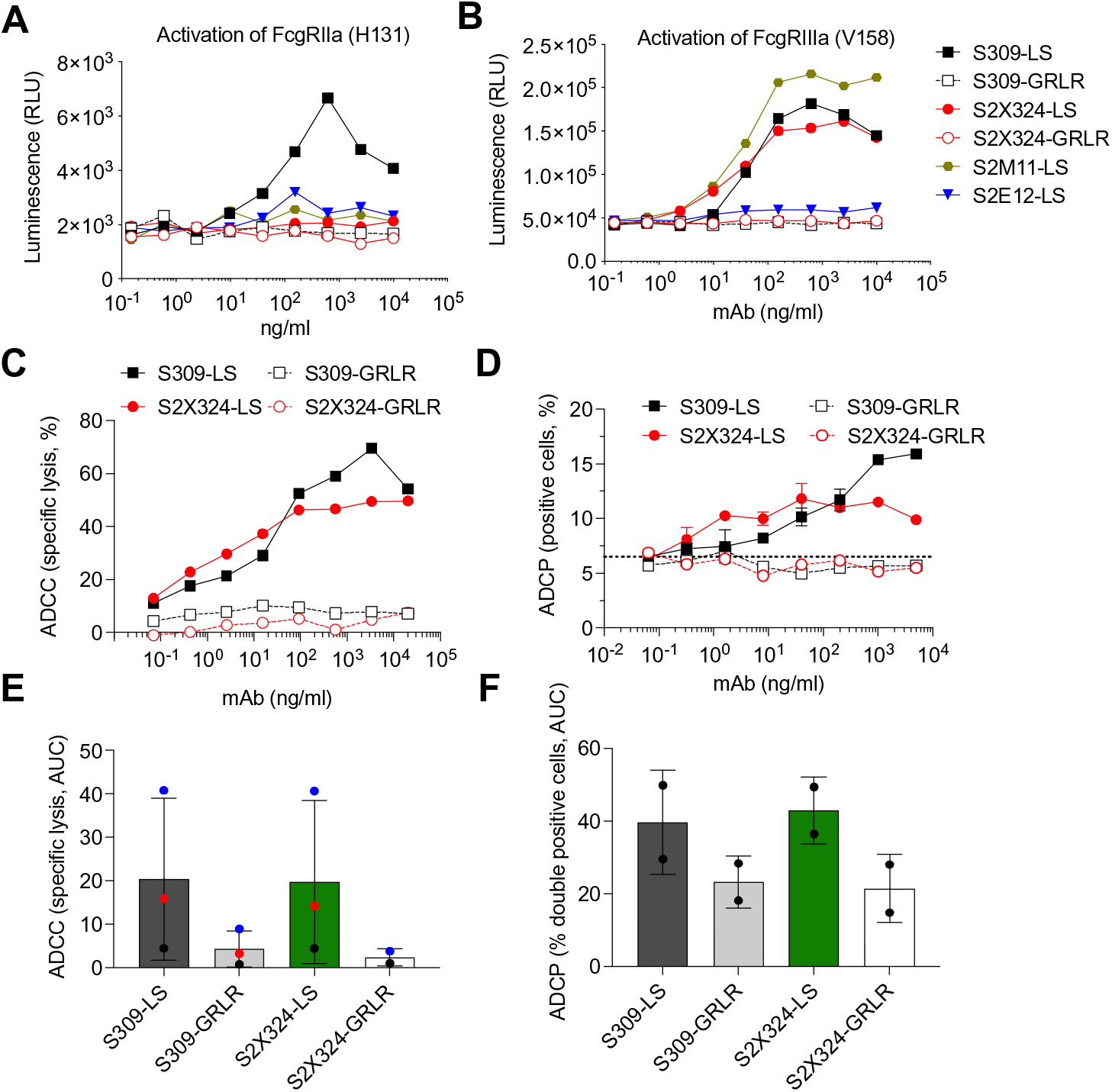
In vitro effector functions by S2X324. **(A-B)**, NFAT-driven luciferase signal induced in Jurkat cells stably expressing FcγRIIa H131 (A) or FcγRIIIa V158 (B) by S2X324 binding to prefusion stabilized SARS-CoV-2 Wuhan S stably expressed in ExpiCHO cells. SE12, S2M11, S309, S309-GRLR, S2X324-GRLR mAbs are included as controls (GRLR, Fc-null). **(C-E)** S2X324-triggered activation of ADCC (C, E) and ADCP (D, F) following incubation with PBMC and monocytes, respectively. AUC analyses of ADCC (**E**) and ADCP (**F**) mediated by S309-LS, S309-GRLR, S2X324-LS and S2X324-GRLR mAbs using primary NK cells or monocytes from three and two donors, respectively.

**Figure S13.**
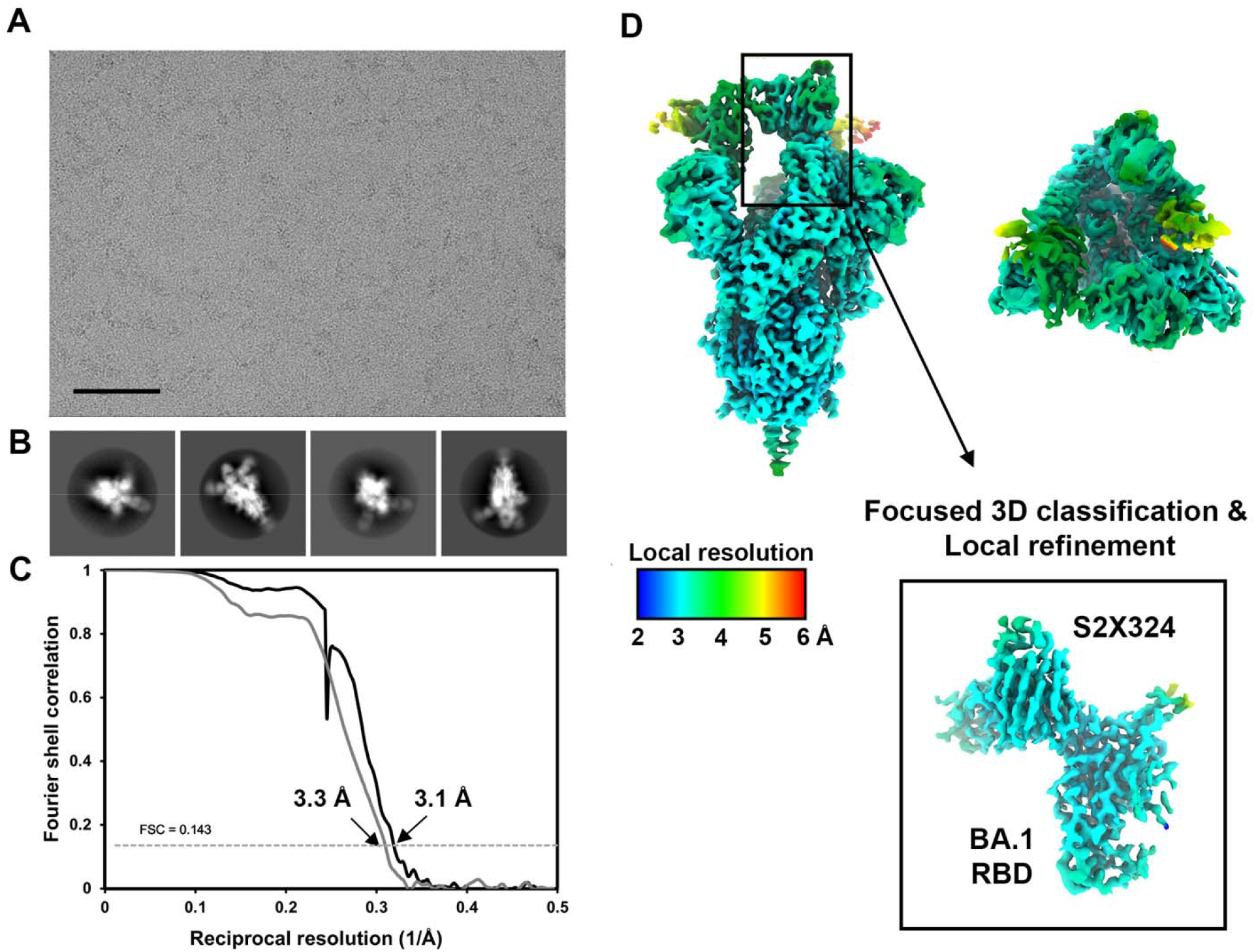
CryoEM data processing of the S2X324-bound SARS-CoV-2 Omicron BA.1 S dataset. **A-B)** Representative electron micrograph (A) and 2D class averages (B) of SARS-CoV-2 Omicron BA.1 S in complex with the S2X324 Fab embedded in vitreous ice. The scale bar represents 100 nm. **C)** Gold-standard Fourier shell correlation curves for the S2X324-bound SARS-CoV-2 S maps with two RBDs open state (black line) and locally refined RBD/S2X324 variable domain (grey line). The 0.143 cutoff is indicated by a horizontal dashed line. **D)** Local resolution maps calculated using CryoSPARC for the SARS-CoV-2 S/S2X324 Fab complex structure in two orthogonal orientations, the side view (left) and the top view (right) as well as for the locally refined RBD/S2X324 variable domain region (inset).

**Figure S14.**
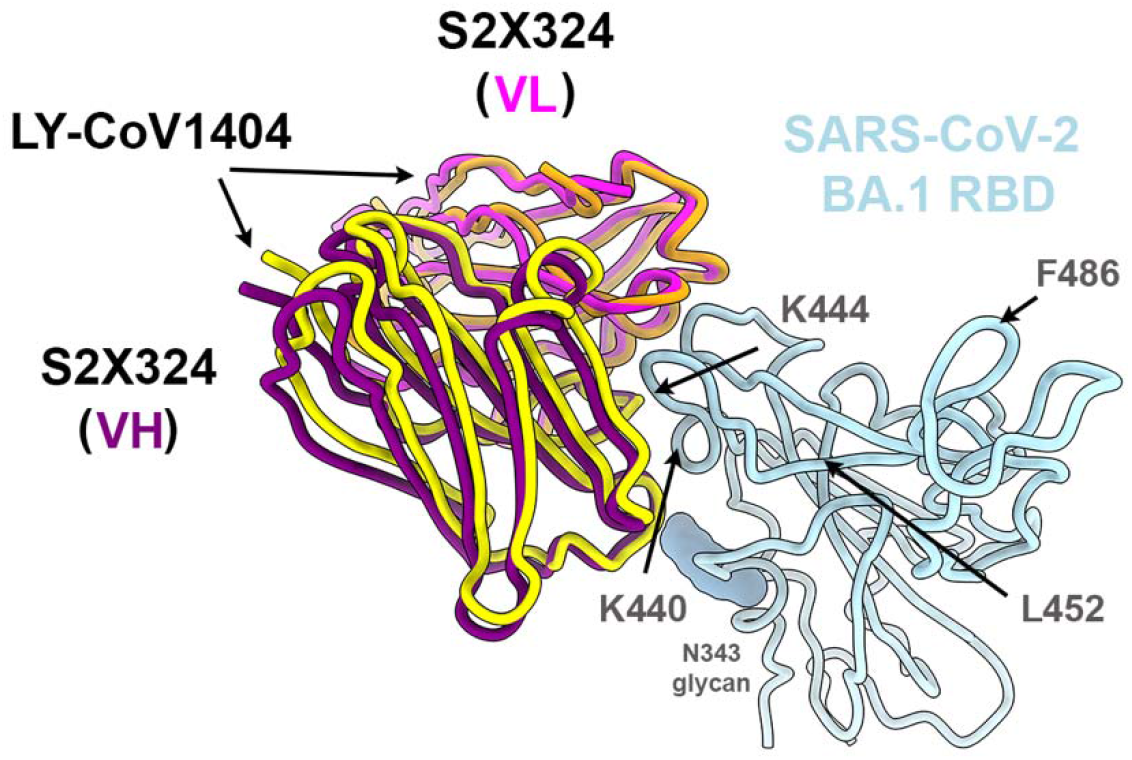
Superposition of S2X324 and LY-CoV1404 with RBD. Ribbon diagram of LY-CoV1404/RBD structure (PDB 7MMO, yellow and orange) superimposed onto S2X324/RBD (CryoEM structure, purple and magenta), using the RBD as a reference. The N343 glycan is rendered as blue spheres. Two selected epitope residues and the BA.5 RBD mutation residues relative to BA.1, L452R and F486V, which are positioned outside the S2X324/LY-CoV1404 epitope are indicated with arrows.

**Fig. S15.**
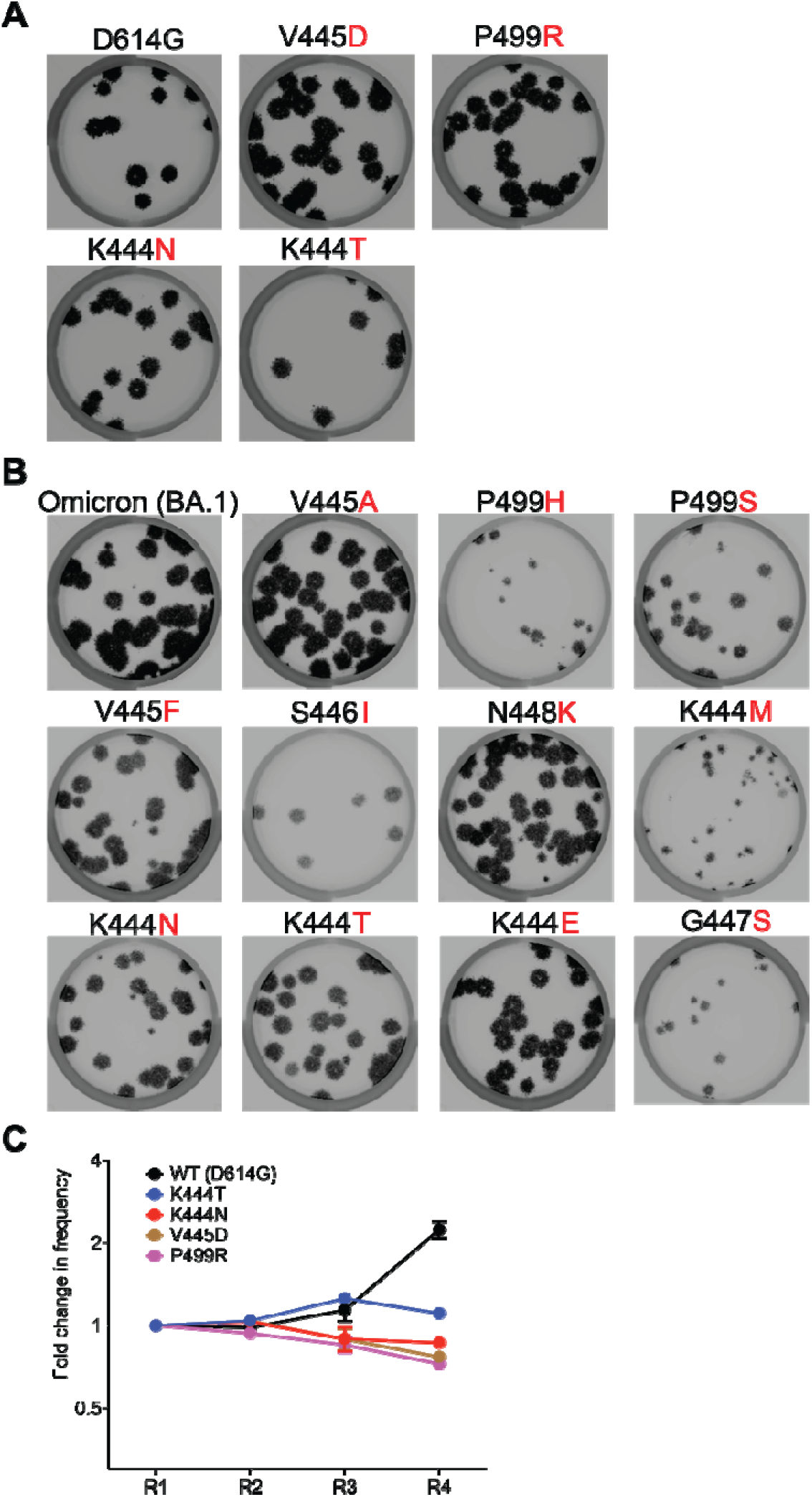
S2X324 escape clone selection by plaque assays using VSV-SARS-CoV-2 Wuhan-Hu-1 D614G and Omicron BA.1 S chimeric viruses. **(A-B)** Plaque assays performed to validate the VSV-SARS-CoV-2 S G614 and Omicron BA.1 mutant in Vero cells in the presence (+) of S2X324 in the overlay. (**C**) Replication fitness of S2X324 escape mutations identified in A on Vero E6 cells.

**Fig. S16.**
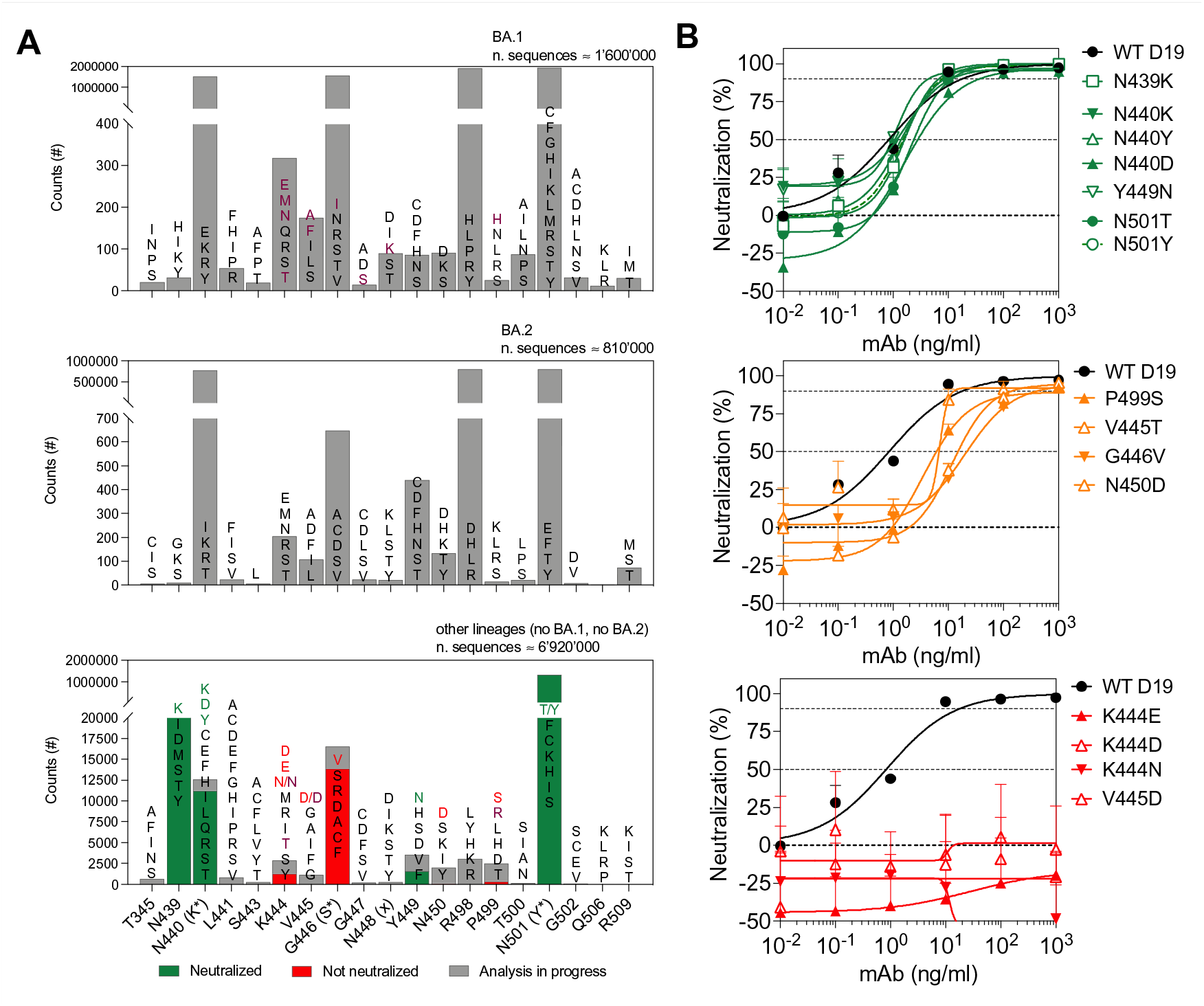
Counts of observed mutants in S2X324 epitope among SARS-CoV-2 sequences. (**A**) Mutants within the S2X324 epitope on the basis of SARS-CoV-2 variant genome sequences available on GISAID as of 11 April 2022. Colored are shown the mutants neutralized (green) or not neutralized (red) by S2X324 and the escape mutants found with VSV chimera (purple). (**B**) Neutralization against VSV pseudoviruses bearing Wuhan-Hu-1/G614 spike with mutations within the epitope of S2X324.

**Table S1.** Neutralizing geometric mean titers (GMTs) against Wu-G614, Delta, BA.1, BA.2, and SARS-CoV S VSV pseudoviruses using plasma from subjects who were infected prior to being vaccinated with 2 doses (n=15) or 3 doses (n=8), subjects who experienced a Delta breakthrough infection with 2 doses (n=15), a BA.1 breakthrough infection with either 2 (n=8) or 3 doses (n=8), or were vaccinated-only with 3 doses (n=7).

| GMTs | Infected-Vaccinated (2 doses) | Infected-Vaccinated (3 doses) | Delta Breakthrough (2 doses) | BA.1 Breakthrough (2 doses) | BA.1 Breakthrough (3 doses) | Vaccinated only (3 doses) |
| --- | --- | --- | --- | --- | --- | --- |
| Wu-G614 | 2065 | 6828 | 2031 | 5388 | 5140 | 1862 |
| Delta | 1398 | 1909 | 1033 | 2778 | 1927 | 662 |
| BA.1 | 576 | 830 | 560 | 4226 | 4608 | 426 |
| BA.2 | 1139 | 1708 | 792 | 2441 | 2464 | 531 |
| SARS-CoV | 129 | 378 | 141 | 158 | 163 | 216 |

**Table S2.** Description of the Cohorts of SARS-CoV-2-immune individuals, Related to Figure 1B-C

| Sample ID | Gender | Vacc. | Doses | COVID diagnosis | Symptom onset | Sample date | $\Delta$ days | Cohort* |
| --- | --- | --- | --- | --- | --- | --- | --- | --- |
| VC202 | F | Yes | 3 | Yes | 2022-03-24 | 2022-04-12 | 19 | 1 |
| VC203 | M | Yes | 3 | Yes | 2022-03-12 | 2022-04-12 | 31 | 1 |
| VC204 | F | Yes | 2 | Yes | 2022-01-19 | 2022-04-12 | 83 | 1 |
| VC206 | F | Yes | 2 | Yes | 2022-03-15 | 2022-04-12 | 28 | 1 |
| VC207 | F | Yes | 3 | Yes | 2022-02-19 | 2022-04-12 | 52 | 1 |
| VC208 | M | Yes | 2 | Yes | 2021-12-31 | 2022-04-12 | 102 | 1 |
| VC209 | F | Yes | 3 | Yes | 2022-03-29 | 2022-04-12 | 14 | 1 |
| VC205 | F | No | 0 | Yes | 2022-01-11 | 2022-04-12 | 91 | 2 |
| VC210 | F | No | 0 | Yes | 2022-01-21 | 2022-04-13 | 82 | 2 |
| VC211 | M | No | 0 | Yes | 2022-03-09 | 2022-04-14 | 36 | 2 |
| VC201 | F | Yes | 3 | No | NA | 2022-04-12 | NA | 3 |
| HCW102 | F | Yes | 2 | Yes | 2020-11-25 | 2021-05-31 | 187 | 3 |
| HCW104 | F | Yes | 2 | Yes | 2020-11-28 | 2021-05-31 | 184 | 3 |
| HCW105 | F | Yes | 2 | Yes | 2020-12-14 | 2021-05-31 | 168 | 3 |
| HCW107 | F | Yes | 2 | Yes | 2020-11-25 | 2021-05-31 | 187 | 3 |
| HCW112 | F | Yes | 2 | Yes | 2020-10-19 | 2021-05-31 | 224 | 3 |
| HCW114 | F | Yes | 2 | Yes | 2020-10-27 | 2021-05-31 | 216 | 3 |
| HCW116 | M | Yes | 2 | Yes | 2020-10-19 | 2021-05-31 | 224 | 3 |
| HCW136 | F | Yes | 2 | Yes | 2020-11-18 | 2021-06-01 | 195 | 3 |
| HCW144 | F | Yes | 2 | Yes | 2020-12-02 | 2021-06-01 | 181 | 3 |
| HCW145 | F | Yes | 2 | Yes | 2020-12-29 | 2021-06-01 | 154 | 3 |

F, female; M, male. \*, 1, Vaccinated and infected with Omicron; 2, Not-vaccinated infected with Omicron; 3, Control pre-Omicron vaccinated

**Table S3.** Neutralizing activity against Wu-G614, Delta, BA.1, BA.2, BA.5 and SARS-CoV S VSV pseudoviruses using plasma from subjects who experienced breakthrough Omicron infection.

|  | Donor ID | Neutralization of VSV-SARS-CoV1/2<br>(reciprocal ID<sub>50</sub>) |  |  |  |  |  | Neutralization of VSV-SARS-CoV1/2<br>(fold loss vs Wu-Hu-1) |  |  |  |  |  | Fold loss of<br>BA.5 vs<br>BA.1 |
| --- | --- | --- | --- | --- | --- | --- | --- | --- | --- | --- | --- | --- | --- | --- |
|  |  | Wu | BA.1 | BA.2 | BA.5 | Delta | SARS-CoV | Wu | BA.1 | BA.2 | BA.5 | Delta | SARS-CoV | BA.5 |
| Vaccinated | VC201 | 1053 | 814 | 229 | 154 | 337 | 114 | 1 | 1 | 5 | 7 | 3 | 9 | 5 |
|  | VC202 | 4208 | 3827 | 2834 | 1467 | 2185 | 162 | 1 | 1 | 1 | 3 | 2 | 26 | 3 |
|  | VC203 | 4706 | 1150 | 1707 | 1416 | 2222 | 264 | 1 | 4 | 3 | 3 | 2 | 18 | 1 |
|  | VC204 | 7028 | 5804 | 2871 | 1085 | 4262 | 160 | 1 | 1 | 2 | 6 | 2 | 44 | 5 |
|  | VC206 | 6901 | 5904 | 2242 | 903 | 2510 | 100 | 1 | 1 | 3 | 8 | 3 | 69 | 7 |
|  | VC207 | 17538 | 1410 | 1533 | 916 | 19167 | 132 | 1 | 12 | 11 | 19 | 1 | 133 | 2 |
|  | VC208 | 6829 | 4333 | 2276 | 635 | 4401 | 126 | 1 | 2 | 3 | 11 | 2 | 54 | 7 |
|  | VC209 | 22780 | 7911 | 6322 | 2590 | 8192 | 368 | 1 | 3 | 4 | 9 | 3 | 62 | 3 |
| Non-vacc. | VC205 | 40 | 564 | 140 | 34 | 23 | 145 | 1 | 0 | 0 | 1 | 2 | 0 | 17 |
|  | VC210 | <10 | 20 | 9 | <10 | <10 | 31 | NC | NC | NC | NC | NC | NC | - |
|  | VC211 | 598 | 7294 | 1556 | 662 | 501 | 86 | 1 | 0 | 0 | 1 | 1 | 7 | 11 |

Shown is the reciprocal of the half-maximum inhibitory dose (ID<sub>50</sub>). NC, not calculable.

**Table S4.**
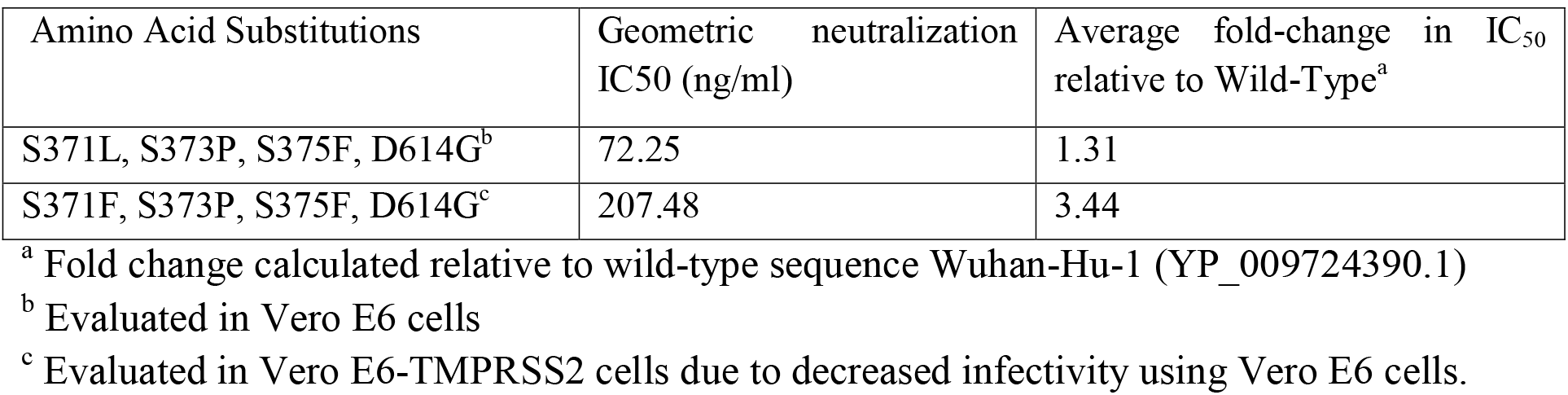
Neutralizing activity of sotrovimab against VSV psuedoviruses harboring the SARS-CoV-2 Wuhan-Hu-1 S (D614) with the indicated substitutions.

**Table S5.**
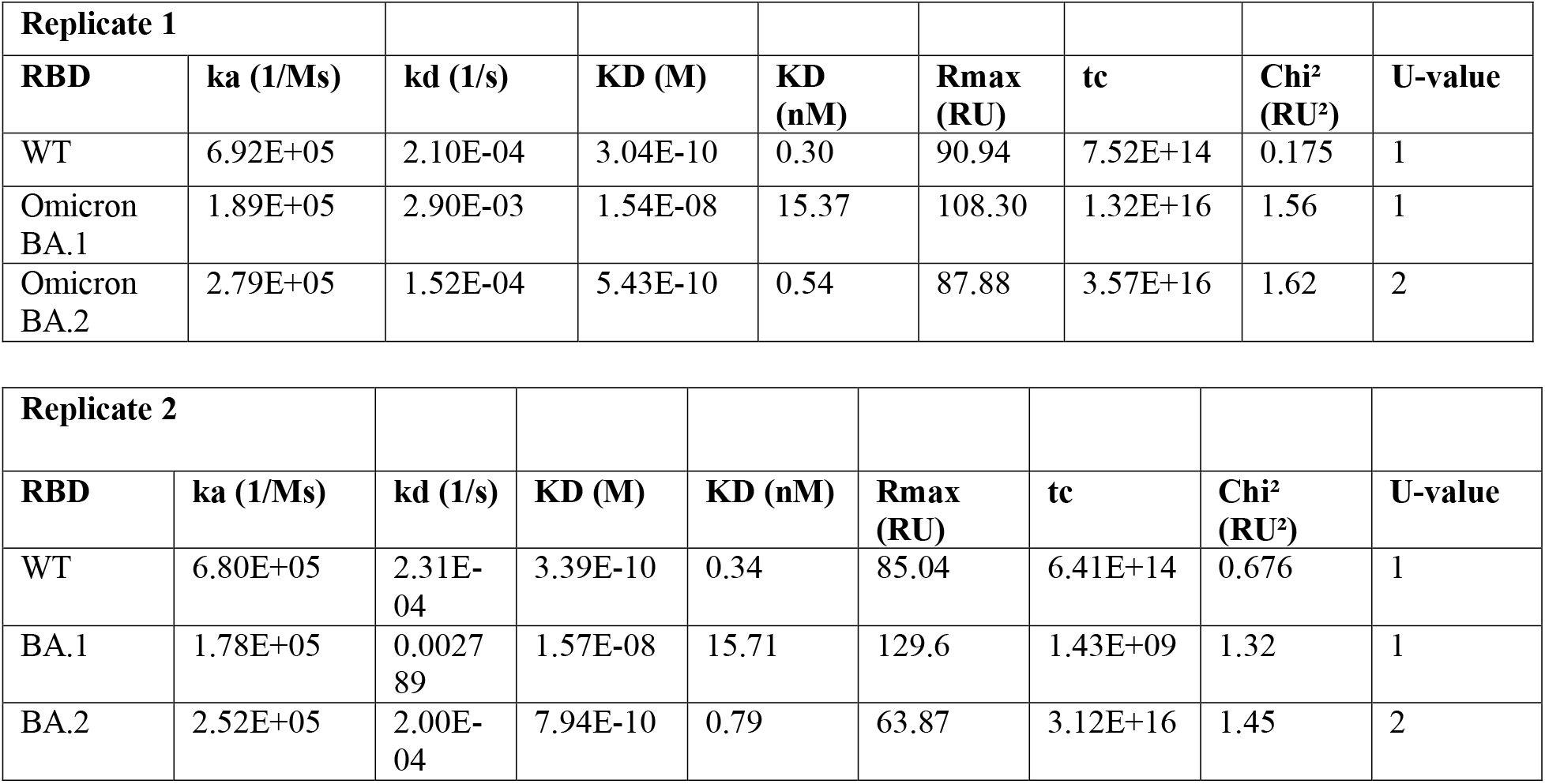
Kinetics parameters of S2X324 Fab binding to different RBDs evaluated from SPR binding assay with 2 replicates.

**Table S6.**
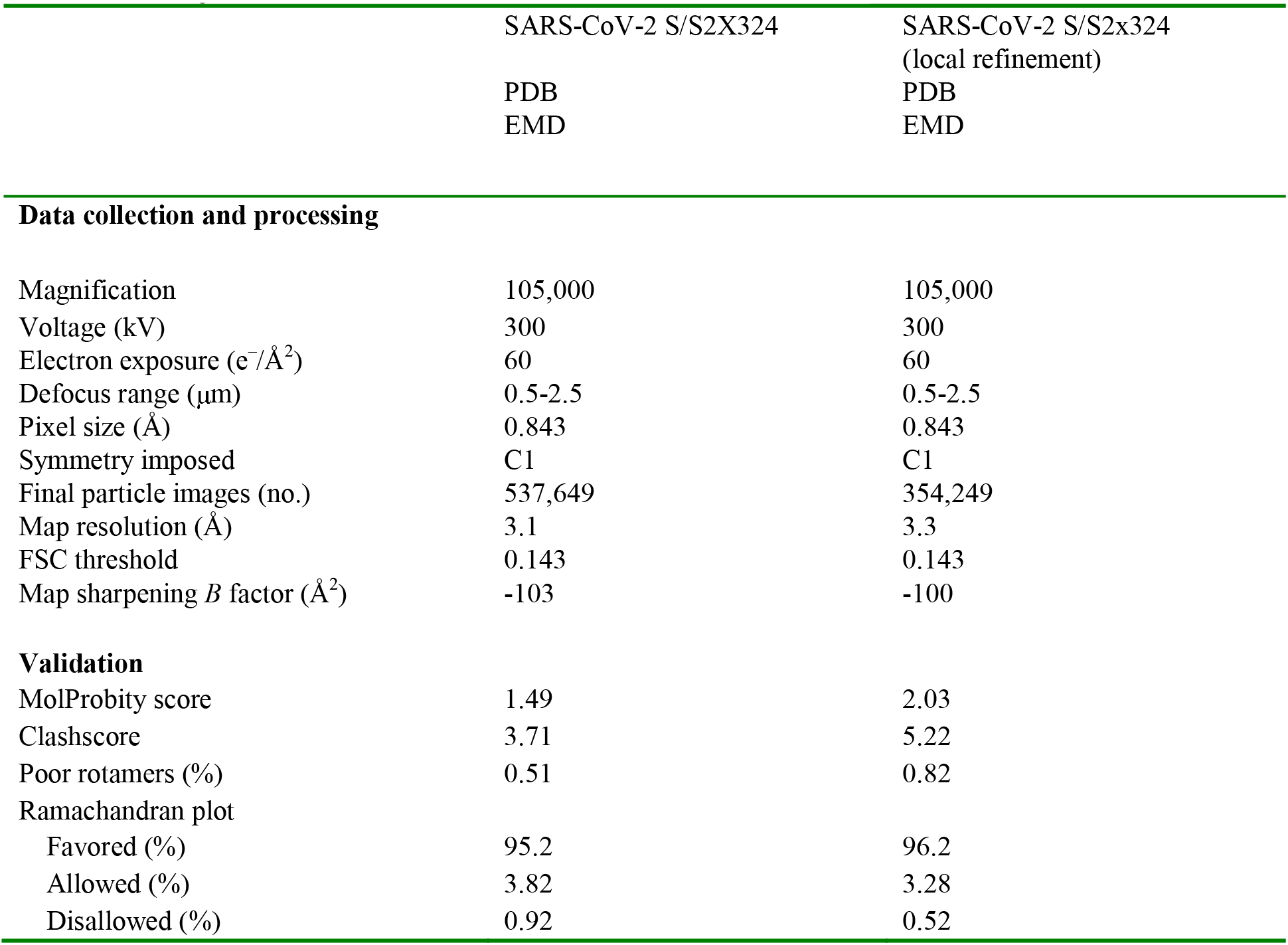
CryoEM data collection and refinement statistics.

|  | SARS-CoV-2 S/S2X324 | SARS-CoV-2 S/S2x324<br>(local refinement) |
| --- | --- | --- |
|  | PDB | PDB |
|  | EMD | EMD |
| **Data collection and processing** |  |  |
| Magnification | 105,000 | 105,000 |
| Voltage (kV) | 300 | 300 |
| Electron exposure (e<sup>-</sup>/Å<sup>2</sup>) | 60 | 60 |
| Defocus range (μm) | 0.5-2.5 | 0.5-2.5 |
| Pixel size (Å) | 0.843 | 0.843 |
| Symmetry imposed | C1 | C1 |
| Final particle images (no.) | 537,649 | 354,249 |
| Map resolution (Å) | 3.1 | 3.3 |
| FSC threshold | 0.143 | 0.143 |
| Map sharpening *B* factor (Å<sup>2</sup>) | -103 | -100 |
| **Validation** |  |  |
| MolProbity score | 1.49 | 2.03 |
| Clashscore | 3.71 | 5.22 |
| Poor rotamers (%) | 0.51 | 0.82 |
| Ramachandran plot |  |  |
| Favored (%) | 95.2 | 96.2 |
| Allowed (%) | 3.82 | 3.28 |
| Disallowed (%) | 0.92 | 0.52 |

**Table S7.**
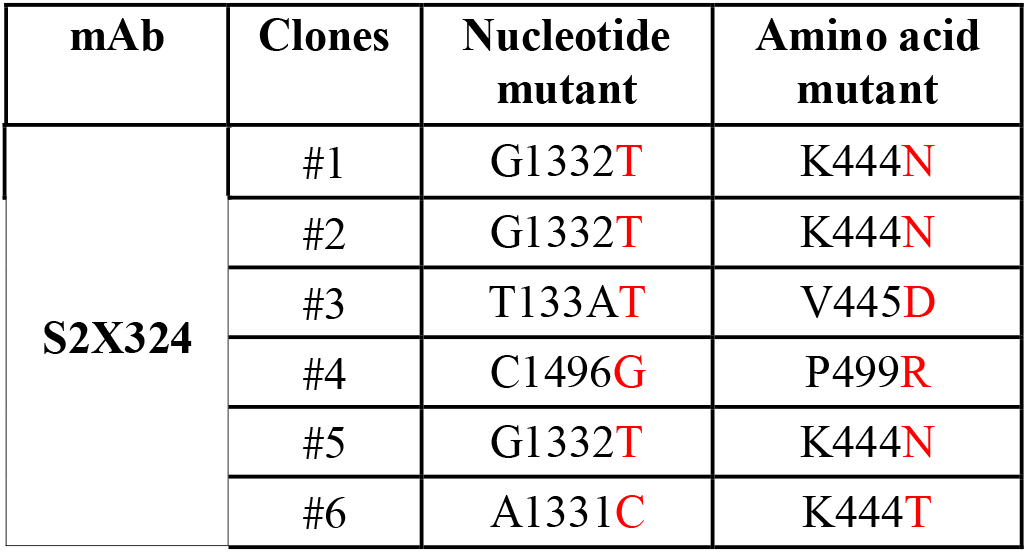
Summary of nucleotide and amino acid mutations found in neutralization-resistant VSV-SARS-CoV-2 G614 S chimera plaques.

| mAb | Clones | Nucleotide mutant | Amino acid mutant |
| --- | --- | --- | --- |
| **S2X324** | #1 | G1332**T** | K444**N** |
|  | #2 | G1332**T** | K444**N** |
|  | #3 | T133A**T** | V445**D** |
|  | #4 | C1496**G** | P499**R** |
|  | #5 | G1332**T** | K444**N** |
|  | #6 | A1331**C** | K444**T** |

**Table S8.**
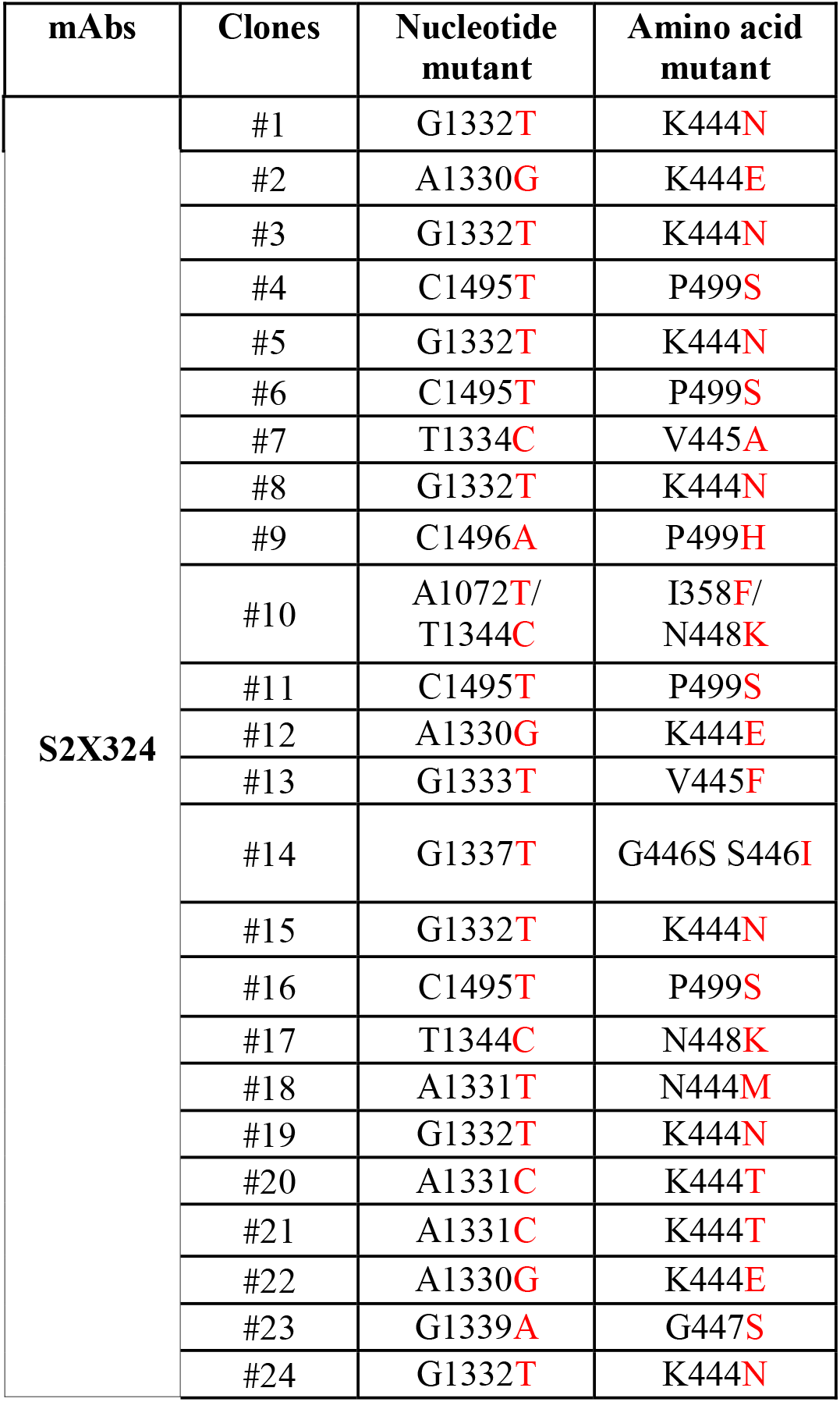
Summary of nucleotide and amino acid mutations found in neutralization-resistant VSV-SARS-CoV-2 Omicron BA.1 S chimera plaques.

**Table S9.** Summary of the mutations retrieved from resistance studies with VSV chimera and their prevalence among Delta and Omicron isolates.

|  | VSV chimera | Delta* | Omicron* | All* |
| --- | --- | --- | --- | --- |
| **K444N** | Wuhan/BA.1 | 950 | 96 | 1,303 |
| **K444T** | Wuhan/BA.1 | 103 | 157 | 265 |
| **K444E** | BA.1 | 20 | 9 | 37 |
| **K444M** | BA.1 | 144 | 10 | 174 |
| **V445A** | BA.1 | 217 | 192 | 613 |
| **V445F** | BA.1 | 246 | 46 | 368 |
| **V445D** | Wuhan | 8 | 40 | 61 |
| **S446I** | BA.1 | 1 | 22 | 23 |
| **G447S** | BA.1 | 34 | 10 | 81 |
| **N448K** | BA.1 | 13 | 3 | 31 |
| **P499H** | BA.1 | 64 | 13 | 116 |
| **P499R** | Wuhan | 1,581 | 12 | 1,643 |
|  |  | 3381 | 610 | 4,715 |
|  |  | 4,146,647 | 3,121,074 | 9,756,845 |
|  |  | 0.082% | 0.020% | 0.048% |

\*Counts retrieved from GISAID as of May 3, 2022, (sequences with Ns>5% were excluded from analysis)

**Table S10.**
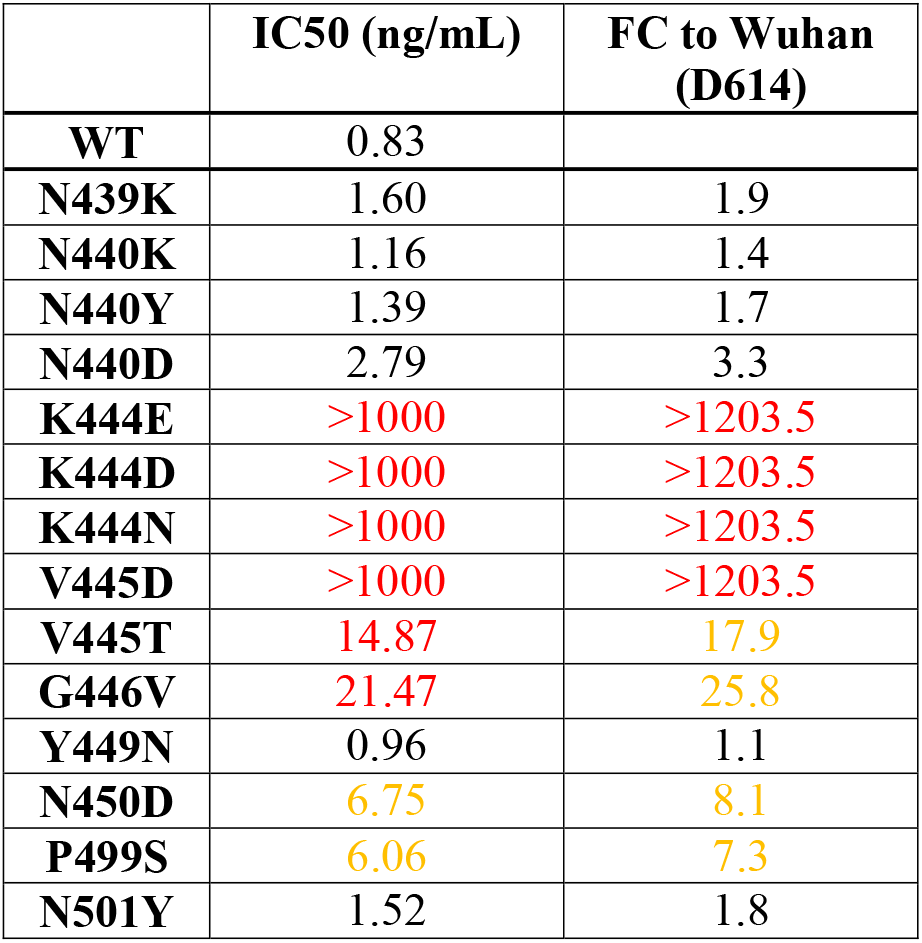
Neutralization against VSV pseudoviruses bearing Wuhan-Hu-1/G614 spike with mutations within the epitope of S2X324.

## Materials and Methods

### Cell lines

Cell lines used in this study were obtained from ATCC (HEK293T and Vero E6), Thermo Fisher Scientific (Expi-CHO-S cells, FreeStyle 293-F cells and Expi293F cells) or Takara (Lenti-X 293T cells). Vero E6 cells stably expressing the TMPRSS2 protease (VeroE6-TMPRSS2 cells) were generated at Vir (*34*). None of the cell lines used was authenticated. Cell lines were routinely tested for mycoplasma contamination.

### Authentic SARS-CoV-2 strains

SARS-CoV-2 isolates used in this study were obtained from BEI (WT: SARS-CoV-2 isolate USA-WA1/2020, BEI ref. NR-52281; BA.1: hCoV-19/USA/MD-HP20874/2021, BEI ref. NR-56461; BA.2 (BEI): hCoV-19/USA/MD-HP24556/2022, BEI ref. NRS-56511; BA.4: hCoV-19/USA/MD-HP30386/2022, BEI ref. NRS-56803; BA.5: hCoV-19/USA/COR-22-063113/2022, BEI ref. NRS-58616); BA.2.12.1: USA/NY-MSHSPSPPV56475/2022, BEI ref. NR-56782 and propagated in house. The genomic sequence of all strains was confirmed by Sanger and NGS sequencing.

### Sample donors

Blood samples were collected from participants as part of the Hospitalized or Ambulatory Adults with Respiratory Viral Infections (HAARVI) study and was approved by the University of Washington Human Subjects Division Institutional Review Board (STUDY00000959). Baseline socio-demographic and clinical data for these individuals are summarized in Data S1.

Samples were obtained from SARS-CoV-2 convalescent and vaccinated individuals under study protocols approved by the local institutional review boards (Canton Ticino Ethics Committee, Switzerland, Comitato Etico Milano Area 1). All donors provided written informed consent for the use of blood and blood derivatives (such as peripheral blood mononuclear cells, sera or plasma) for research.

### Viral sequencing for Breakthrough cases

Participants were enrolled after SARS-CoV-2 positive RT-PCR results that showed S-gene dropout or delay using a primer set that predates the S probe set in the “TaqPath” test kit sold by Thermo Fisher. Specifically, the Delta S 157-158 deletion causes S-gene dropout in our diagnostic RT-qPCR assay. The RT-qPCR assay uses probes for human RNase P and SARS-CoV-2 Orf1b and S gene. S-gene dropout or delay was identified using the following individual sample criteria: 1) internal control RNase P amplification was present, 2) Orf1b amplification was present, and 3) S-gene specific probes showed no amplification or a delay of >5 cycles compared to Orf1b due to the 156-157 deletion in delta. Three samples (278C, 284C, and 286C) included in this analysis were either not sequenced due to low volumes or failed sequencing due to low viral loads, but according to outbreaks.info over 90% of all cases in Washington state at the time were Delta SARS-CoV-2.

SARS-CoV-2 genome sequencing was then conducted using a targeted enrichment approach. SARS-CoV-2 was detected using a laboratory-developed test (LDT) or research assay. For the LDT, SARS-CoV-2 detection was performed using real-time RT-PCR with a probe sets targeting Orf1b and S with FAM fluor (Life Technologies 4332079 assays # APGZJKF and #APXGVC4) multiplexed with an RNaseP probe set with VIC or HEX fluor (Life Technologies A30064 or IDT custom) each in duplicate on a QuantStudio 6 instrument (Applied Biosystems). RNA from positive specimens was converted to cDNA using random hexamers and reverse transcriptase (Superscript IV, Thermo) and a sequencing library was constructed using the Illumina TruSeq RNA Library Prep for Enrichment kit (Illumina). The sequencing library was enriched for SARS-CoV-2 using the Twist Respiratory Virus Research Panel (Twist). Libraries were sequenced on a MiSeq or NextSeq instruments. The resulting reads were assembled against the SARS-CoV-2 reference genome Wuhan-Hu-1/2019 (Genbank accession MN908947) using the bioinformatics pipeline https://github.com/seattleflu/assembly. Consensus sequences were deposited to Genbank and GISAID (*80*). Sequence quality was determined using Nextclade version 1.0.0-alpha.8 (https://clades.nextstrain.org/). Lineage was assigned using the Pangolin COVID-19 Lineage Assigner version 3.1.11 (https://pangolin.cog-uk.io/).

### Recombinant Protein Expression and purification for Nasal Swab ELISAs

The SARS-CoV-2 VFLIP ectodomain (*81*) was produced in Expi293F Cells (ThermoFisher Scientific) grown in suspension using Expi293 Expression Medium (ThermoFisher Scientific) at 37°C in a humidified 8% CO_2_ incubator with constant rotation at 130 rpm. Cells grown to a density of 3 million cells per mL were transfected using the ExpiFectamine 293 Transfection Kit (ThermoFisher Scientific) and cultivated for four days prior to supernatent harvest. SARS-CoV-2 VFLIP was purified from clarified supernatants using a HisTrapHP column (Cytiva) and washed with 10 column volumes of 25 mM sodium phosphate pH 8.0 and 150 mM NaCl before elution on a gradient up to 500 mM imidazole. Purified protein was buffer exchanged into 20 mM Tris-HCl pH 8.0 and 100 mM NaCl, concentrated using 100 kDa MWCO centrifugal filters (Amicon Ultra) to 1-2 mg/mL, and flash frozen.

### AMBRA (antigen-specific memory B cell repertoire analysis) of secreted IgGs

Replicate cultures of total unfractionated PBMC from SARS-CoV-2 infected or vaccinated individuals were seeded in 96 U-bottom plates (Corning) in RPMI1640 supplemented with 10% Hyclone, sodium pyruvate, MEM non-essential amino acid, stable glutamine and Penicillin-Streptomycin. Memory B cell stimulation and differentiation was induced by adding 2.5 μg/ml R848 (3 M) and 1000 U/ ml human recombinant IL-2 at 37 °C and 5% CO2. After 10 days, the cell culture supernatants were collected for further analysis by ELISA.

### Analysis of IgGs secreted by plasma cells

Plasma cells from peripheral blood were stained with PE-conjugated anti-CD138 (BD-Pharmingen), enriched by magnetic separation with anti-PE microbeads (Miltenyi) and finally purified by cell sorting on Sony cell sorter. Cells were seeded at 0.5 cells/well in multiple 384 well plates in 50 μl complete RPMI 1640 medium supplemented with 10% FCS (Hyclone) and 10 ng/ml human r-IL6 (R&D). After three days 20 μl of culture supernatants were collected using an automated liquid handling equipment (Perkin Elmer) and tested in parallel micro-ELISA (5 μl per test) for the presence of IgG antibodies against different SARS-CoV-2 RBDs and PBS as negative control.

#### Generation of VSV pseudovirus

Replication defective VSV pseudovirus expressing SARS-CoV-2 spike proteins corresponding to the ancestral Wuhan-Hu-1 virus and the VOCs were generated as previously described (*3*), with some modifications. Lenti-X 293T cells (Takara) were seeded in 15-cm2 dishes at a density of 10L×L106 cells per dish and the following day were transfected with 25 µg of spike expression plasmid with TransIT-Lenti (Mirus, 6600) according to the manufacturer’s instructions. One day after transfection, cells were infected with VSV-luc (VSV-G) with a multiplicity of infection (MOI) of 3 for 1 h, rinsed three times with PBS containing Ca2+ and Mg2+, then incubated for an additional 24 h in complete medium at 37 °C. The cell supernatant was clarified by centrifugation, aliquoted, and frozen at −80 °C. For pseudoviruses expressing S substitutions that resulted in decreased infectivity using Vero E6 cells, Vero E6-TMPRSS2 cells were substituted as target cells to determine pseudovirus titer. Spike expression plasmids used for the generation of VSV pseudoviruses carry the following mutations: <u>BA.1</u>: A67V, Δ69-70, T95I, G142D, Δ143-145, Δ211, L212I, ins214EPE, G339D, S371L, S373P, S375F, K417N, N440K, G446S, S477N, T478K, E484A, Q493R, G496S, Q498R, N501Y, Y505H, T547K, D614G, H655Y, N679K, P681H, N764K, D796Y, N856K, Q954H, N969K, L981F; <u>BA.2</u>: T19I, L24-, P25-, P26-, A27S, G142D, V213G, G339D, S371L, S373P, S375F, D405N, R408S, K417N, N440K, S477N, T478K, E484A, Q493R, Q498R, N501Y, Y505H, D614G, H655Y, N679K, P681H, N764K, D796Y/ N856K/ Q954H/ N969K; <u>BA.3</u>: A67V, H69-, V70-, T95I, G142D, V143-, Y144-, Y145-, N211-, L212I, G339D, S371F, S373P S375F, D405N, K417N, N440K, G446S, S477N, T478K, E484A, Q493R, Q498R, N501Y, Y505H, D614G, H655Y, N679K, P681H, N764K, D796Y, Q954H, N969K; <u>BA.4-N658S</u>: T19I, L24-, P25-, P26-, A27S, G142D, V213G, Δ69-70, G339D, S371F, S373P, S375F, T376A, D405N, R408S, K417N, N440K, L452R, S477N, T478K, E484A, F486V, Q498R, N501Y, Y505H, D614G, H655Y, N658S, N679K, P681H, N764K, D796Y, Q954H, N969K; <u>BA.4-V3G</u>: V3G, T19I, L24-, P25-, P26-, A27S, G142D, V213G, Δ69-70, G339D, S371F, S373P, S375F, T376A, D405N, R408S, K417N, N440K, L452R, S477N, T478K, E484A, F486V, Q498R, N501Y, Y505H, D614G, H655Y, N679K, P681H, N764K, D796Y, Q954H, N969K; <u>BA.5</u>: T19I, L24-, P25-, P26-, A27S, G142D, V213G, Δ69-70, G339D, S371F, S373P, S375F, T376A, D405N, R408S, K417N, N440K, L452R, S477N, T478K, E484A, F486V, Q498R, N501Y, Y505H, D614G, H655Y, N679K, P681H, N764K, D796Y, Q954H, N969K; <u>BA.2.12.1</u>: T19I, L24-, P25-, P26-, A27S, G142D, V213G, G339D, S371F, S373P, S375F, T376A, D405N, R408S, K417N, N440K, L452Q, S477N, T478K, E484A, Q493R, Q498R, N501Y, Y505H, D614G, H655Y, N679K, P681H, S704L, N764K, D796Y, Q954H, N969K; BA.2.75: T19I, del24-26, A27S, G142D, K147E, W152R, F157L, I210V, V213G, G257S, G339H, S371F, S373P, S375F, T376A, D405N, R408S, K417N, N440K, G446S, N460K, S477N, T478K, E484A, Q498R, N501Y, Y505H, D614G, H655Y, N679K, P681H, N764K, D796Y, Q954H, N969K.

#### VSV pseudovirus neutralization

Vero E6 cells were grown in DMEM supplemented with 10% FBS and seeded into clear bottom white 96 well plates (PerkinElmer, 6005688) at a density of 20,000 cells per well. The next day, monoclonal antibodies were serially diluted in pre-warmed complete medium, mixed with pseudoviruses and incubated for 1 h at 37 °C in round bottom polypropylene plates. Medium from cells was aspirated and 50 µl of virus–monoclonal antibody complexes were added to cells, which were then incubated for 1 h at 37 °C. An additional 100 µl of pre-warmed complete medium was then added on top of complexes and cells were incubated for an additional 16–24 h. Conditions were tested in duplicate wells on each plate and eight wells per plate contained untreated infected cells (defining the 0% of neutralization, ‘MAX RLU’ value) and infected cells in the presence of S309 and S2X259 at 20LµgLml−1 each (defining the 100% of neutralization, ‘MIN RLU’ value). For sotrovimab experiments, conditions were tested in triplicate wells starting at 10 µgLml−1 sotrovimab; cells only and untreated infected cells were included on each plate. Virus–monoclonal antibody-containing medium was then aspirated from cells and 100 µl of a 1:2 dilution of SteadyLite Plus (PerkinElmer, 6066759) in PBS with Ca2+ and Mg2+ was added to cells. Plates were incubated for 15 min at room temperature and then analyzed on the Synergy-H1 (Biotek). The average relative light units (RLUs) of untreated infected wells (MAX RLUave) wereas subtracted by the average of MIN RLU (MIN RLUave) and used to normalize percentage of neutralization of individual RLU values of experimental data according to the following formula: (1L−L(RLUxL–LMINLRLUave)/ (MAXLRLUaveL–LMINLRLUave))L×L100. Data were analyzed with Prism (v.9.1.0). IC50 values were calculated from the interpolated value from the log(inhibitor) versus response, using variable slope (four parameters) nonlinear regression with an upper constraint of ≤100, and a lower constrain equal to 0. Each neutralization experiment was conducted on two independent experiments – that is, biological replicates – in which each biological replicate contains a technical duplicate or triplicate. IC50 values across biological replicates are presented as geometric meanL±Ls.d. For sotrovimab, IC50 values were calculated as described above using an upper constraint of <100. For pseudoviruses expressing spike substitutions that resulted in decreased infectivity using Vero E6 cells, Vero E6-TMPRSS2 cells were substituted as target cells for neutralization assays.

### Neutralization of authentic SARS-CoV-2 viruses

Vero-TMPRSS2 cells were seeded into black-walled, clear-bottom 96-well plates at 2×10^4^ cells/well and cultured overnight at 37°C. The next day, 9-point 4-fold serial dilutions of mAbs were prepared in growth media (DMEM + 10% FBS). The different SARS-CoV-2 strains were diluted in infection media (DMEM + 2% BSA) at a final MOI of 0.01 PFU/cell, added to the mAb dilutions and incubated for 30 min at 37°C. Media was removed from the cells, mAb-virus complexes were added and incubated at 37°C for 18h (WT), 21±3h (BA.4, BA.5), 24h (BA.2, BA.2.12.1) or 30h (BA.1). Cells were fixed with 4% PFA (Electron Microscopy Sciences, #15714S), permeabilized with Triton X-100 (SIGMA, #X100-500ML) and stained with an antibody against the viral nucleocapsid protein (Sino Biologicals, #40143-R001) followed by a staining with the nuclear dye Hoechst 33342 (Fisher Scientific, # H1399) and a goat anti-rabbit Alexa Fluor 647 antibody (Invitrogen, #A-21245). Plates were imaged on a Cytation5 plate reader. Whole well images were acquired (12 images at 4X magnification per well) and nucleocapsid-positive cells were counted using the manufacturer’s software.

### Enzyme-linked immunosorbent assay (ELISA)

Ninety-six half area well-plates (Corning, 3690) were coated overnight at 4 °C with 25 μl of sarbecovirus RBD proteins of SARS-CoV-2 (YP_009724390.1), RaTG13 (QHR63300.2), Pangolin_Guangdong-2019 (EPI_ISL_410721), Pangolin_Guangxi, SARS-CoV Urbani (AAP13441.1) and WIV1 (AGZ48831.1), prepared at 5 μg ml−1 in PBS pH 7.2. Plates were then blocked with PBS 1% BSA (Sigma-Aldrich, A3059) and subsequently incubated with mAb serial dilutions for 1 h at room temperature. After 4 washing steps with PBS 0.05% Tween 20 (PBS-T) (Sigma-Aldrich, 93773), goat anti-human IgG secondary antibody (Southern Biotech, 2040-04) was added and incubated for 1 h at room temperature. Plates were then washed four times with PBS-T and 4-nitrophenyl phosphate (pNPP, Sigma-Aldrich, 71768) substrate was added. After 30 min incubation, absorbance at 405 nm was measured by a plate reader (Biotek) and data were plotted using Prism GraphPad 9.1.0.

### Nasal Swab ELISA

For anti-S ELISA, 50 μL of 2 μg/mL S was plated onto 384-well Nunc Maxisorp (ThermoFisher) plates in PBS and sealed overnight at room temperature. The next day plates were washed 4 × in Tris Buffered Saline Tween (TBST-20mM Tris pH 8, 150mM NaCl, 0.1% Tween) using a plate washer (BioTek) and blocked with Casein (ThermoFisher) for 1 h at 37°C. Plates were washed 4 × in TBST and 1:5 serial dilutions of nasal swab in universal transport media were made in 50 μL TBST and incubated at 37°C for 1 h. Plates were washed 4 × in TBST, then anti-human (Invitrogen) horseradish peroxidase-conjugated antibodies were diluted 1:5,000 and 50 μL added to each well and incubated at 37°C for 1 h. Plates were washed 4 × in TBST and 50 μL of TMB (SeraCare) was added to every well for 3 min at room temperature. The reaction was quenched with the addition of 50 μL of 1 N HCl. Plates were immediately read at 450 nm on a VarioSkanLux plate reader (ThermoFisher) and data plotted and fit in Prism (GraphPad) using nonlinear regression sigmoidal, 4PL, X is log(concentration) to determine EC_50_ values from curve fits. Where the curve did not reach an OD450 of 4, a constraint of OD450 4 was placed on the upper bounds of the fit.

### Recombinant protein production for SPR binding assays

SARS-CoV-2 RBD constructs contain residues 328–531 of the spike protein from GenBank NC_045512.2 with an N-terminal signal peptide and a C-terminal thrombin cleavage site-Twin-Strep-8×His-tag. For SPR binding assays, proteins were expressed in Expi293F cells (Thermo Fisher Scientific) at 37 °C and 8% CO2. Transfections were performed using the ExpiFectamine 293 Transfection Kit (Thermo Fisher Scientific). Cell culture supernatants were collected three to five days after transfection and supplemented with 10x PBS to a final concentration of 2.5x PBS (342.5 mM NaCl, 6.75 mM KCl and 29.75 mM phosphates). SARS-CoV-2 RBDs were purified using a Cobalt affinity column (HisTALON Superflow column from Takara or HiTrap TALON crude column from Cytiva) followed by buffer exchange into PBS using a HiPrep 26/10 desalting column (Cytiva) or, for the Omicron BA.1 and BA.2 RBDs, a Superdex 200 Increase 10/300 GL column (Cytiva).

### Surface plasmon resonance (SPR) binding assays to measure binding of RBDs with S2X324 Fab

Measurements were performed using a Biacore T200 instrument. A CM5 chip with covalently immobilized StrepTactin XT was used for surface capture of Twin-Strep Tag containing RBDs. Running buffer was HBS-EP+ pH 7.4 (Cytiva) and measurements were performed at 25 LJC. Experiments were performed with a 3-fold dilution series of monomeric S2X324 Fab: 300, 100, 33, 11 nM and were run as single-cycle kinetics. Data were double reference-subtracted and fit to a binding model using Biacore Evaluation software. The 1:1 binding model was used to estimate the kinetics parameters. The experiment was performed twice with two biological replicates for each ligand (RBDs). KD values were reported as the average of two replicates with the corresponding standard deviation.

### Competition assay by biolayer interferometry

Biolayer interferometry was used to assess S2X324 competition with S2K146 and S309 using an Octet HTX (Sartorius). All reagents were prepared in kinetics buffer (PBS 0.01% BSA) at the indicated concentrations. His-tagged SARS-CoV-2 RBD was prepared at 8 μg ml−1 and loaded on prehydrated anti-penta-HIS biosensors (Sartorius) for 3 min. Biosensors were then moved into a solution containing S2X324 mAb (20 μg ml−1) and association recorded for 7 min. A second association step was subsequently performed into S2X324 (as control), S2K146 and S309 mAbs solutions at 20 μg ml−1 and recorded for 7 min. Response values were exported and plotted using GraphPad Prism 9.1.0.

### Blockade of SARS-CoV-2 binding to ACE2

SARS-CoV-2 mouse/rabbit Fc-tagged RBDs (final concentration 20 ng/ml) were incubated with serially diluted recombinant mAbs (from 25 µg/ml) and incubated for 1 h 37°C. The complex RBD:mAbs was then added to a pre-coated hACE2 (2 µg/ml in PBS) 96-well plate MaxiSorp (Nunc) and incubated 1 hour at room temperature. Subsequently, the plates were washed, and a goat anti-mouse/rabbit IgG (Southern Biotech) coupled to alkaline phosphatase (Jackson Immunoresearch) added to detect mouse Fc-tagged RBDs binding. After further washing, the substrate (p-NPP, Sigma) was added, and plates read at 405 nm using a microplate reader (Biotek). The percentage of inhibition was calculated as follow: (1-((OD sample-OD neg ctr)/(OD pos. ctr-OD neg. ctr))*100

### Cell-surface mAb-mediated S_1_ shedding

CHO cells stably expressing the prototypic SARS-CoV-2 S were harvested, washed in wash buffer (PBS 1% BSA 2 mM EDTA) and resuspended in PBS. Cells were then counted and 90’000 cells/well were dispensed into a round-bottom 96 well plate (Corning) to be treated with 10 ug/ml TPCK-Trypsin (Worthington Biochem) for 30 min at 37°C. After a washing step, cells were incubated with 15 µg/ml mAbs solution for 180, 120, 60, 30 or 5 min at 37°C. After the incubation for the allotted time, cells were washed with ice-cold wash buffer and stained with 1.5 µg/ml Alexa Fluor647-labeled Goat Anti-Human IgG secondary Ab (Jackson Immunoresearch) for 30 min on ice in the dark. Cells were then washed twice with cold wash buffer and analyzed using a ZE5 cytometer (Biorad) with acquisition chamber T= 4°C. Binding at each time point (MFI) was determined normalizing to the MFI at 5 minutes time point and data plotted using GraphPad Prism v. 9.1.1

### CryoEM sample preparation, data collection and data processing

Recombinantly expressed and purified S2X324 Fab and SARS-CoV-2 Omicron BA.1 S (*17*) were incubated at 1 mg/ml (for UltraAuFoil grids) or 0.1mg/ml (for lacey thin carbon grids) with a 1.2 molar excess of S2X324 Fab at 4°C for 1 hr. Three microliters of the complex mixture were loaded onto freshly glow discharged R 2/2 UltrAuFoil grids (200 mesh) or lacey grids covered with a thin layer of home-made carbon, prior to plunge freezing using a Vitrobot MarkIV (ThermoFisher Scientific) with a blot force of 0 and 6 sec blot time (for the UltrAuFoil grids) or with a blot force of -1 and 3 sec blot time (for the lacey thin carbon grids) at 100 % humidity and 22°C.

Data were acquired using an FEI Titan Krios transmission electron microscope operated at 300 kV and equipped with a Gatan K3 direct detector and Gatan Quantum GIF energy filter, operated in zero-loss mode with a slit width of 20 eV. Automated data collection was carried out using Leginon (*82*) at a nominal magnification of 105,000x with a pixel size of 0.843 Å. The dose rate was adjusted to 15 counts/pixel/s, and each movie was acquired in super-resolution mode fractionated in 75 frames of 40 ms. 10,842 micrographs were collected with a defocus range comprised between -0.5 and -2.5 μm. Movie frame alignment, estimation of the microscope contrast-transfer function parameters, particle picking, and extraction were carried out using Warp (*83*).

Two rounds of reference-free 2D classification were performed using CryoSPARC (*84*) to select well-defined particle images. These selected particles were subjected to two rounds of 3D classification with 50 iterations each (angular sampling 7.5° for 25 iterations and 1.8° with local search for 25 iterations), using our previously reported closed SARS-CoV-2 S structure as initial model (PDB 6VXX) (*85*) using Relion (*86*). 3D refinements were carried out using non-uniform refinement along with per-particle defocus refinement in CryoSPARC (*87*). Selected particle images were subjected to the Bayesian polishing procedure (*88*) implemented in Relion3.0 before performing another round of non-uniform refinement in CryoSPARC followed by per-particle defocus refinement and again non-uniform refinement. To improve the density of the S/S2X324 interface, the particles from the non-uniform refinement were subjected to focus 3D classification without refining angles and shifts using a soft mask on the closed RBD and bound S2X324 variable domains with a tau value of 60 in Relion. Particles belonging to classes with the best resolved local density were selected and subject to local refinement using CryoSPARC. Local resolution estimation, filtering, and sharpening were carried out using CryoSPARC. Reported resolutions are based on the gold-standard Fourier shell correlation (FSC) of 0.143 criterion and Fourier shell correlation curves were corrected for the effects of soft masking by high-resolution noise substitution (*89, 90*).

### Model building and refinement

UCSF Chimera (*91*) and Coot (*92*) were used to fit atomic models into the cryoEM maps. The Omicron BA.1 S/S2X324 and RBD/S2X324 models were refined and relaxed using Rosetta using sharpened and unsharpened maps (*93, 94*).

### Selection of SARS-CoV-2 monoclonal antibody escape mutants

A VSV-SARS-CoV-2 Wuhan-Hu-1 D614G S and Omicron BA.1 S chimera were used to select for mAb resistant mutants, as previously described (*35*). Briefly, mutants were recovered by plaque isolation on Vero E6 cells with the indicated mAb in the overlay. The concentration of mAb in the overlay was determined by neutralization assays at a multiplicity of infection (MOI) of 100. Escape clones were plaque-purified on Vero E6 cells in the presence of mAb, and plaques in agarose plugs were amplified on MA104 cells with the mAb present in the medium. Viral stocks were amplified on MA104 cells at an MOI of 0.01 in Medium 199 containing 2% FBS and 20 mM HEPES pH 7.7 (Millipore Sigma) at 34°C. Viral supernatants were harvested upon extensive cytopathic effect and clarified of cell debris by centrifugation at 1,000 x g for 5 min. Aliquots were maintained at -80°C. Viral RNA was extracted from VSV-SARS-CoV-2 S mutant viruses using RNeasy Mini kit (Qiagen), and the S gene was amplified using OneStep RT-PCR Kit (Qiagen). The mutations were identified by Sanger sequencing (GENEWIZ). Their resistance was verified by subsequent virus infection in the presence or absence of mAb. Vero E6 cells were seeded into 12 well plates overnight. The virus was serially diluted using DMEM and cells were infected at 37°C for 1 h. Cells were cultured with an agarose overlay in the presence or absence of mAb at 34°C for 2 days. Plates were scanned on a biomolecular imager and expression of eGFP monitored at 48 hours post-infection.

### Viral replication fitness assays

Vero E6 cells were seeded at 1×106 cells per well in 6-well plates. Cells were infected with MOI of 0.02, with VSV-SARS-CoV-2 Wuhan-Hu-1 D614G and four escape mutants mixed at equal (0.20) frequencies. Following 1 h incubation, cell monolayers were washed three times with HBBS and cultures were incubated for 72 h in humidified incubators at 34°C. To passage the progeny viruses, virus mixture was continuously passaged four times in Vero E6 cells at MOI of 0.02. Cellular RNA samples from each passage were extracted using RNeasy Mini kit (QIAGEN) and subjected to next-generation sequencing as described previously to confirm the introduction and frequency of substitutions (*45*).

### Measurement of Fc-effector functions

S2X324-dependent activation of human FcγRIIa and IIIa was performed with a bioluminescent reporter assay. ExpiCHO cells stably expressing full-length wild-type SARS-CoV-2 S (target cells) were incubated with different amounts of mAbs. After a 25-minute incubation, Jurkat cells stably expressing FcγRIIIa receptor (V158 variant) or FcγRIIa receptor (H131 variant) and NFAT-driven luciferase gene (effector cells) were added at an effector to target ratio of 6:1 for FcγRIIIa and 5:1 for FcγRIIa. Signaling was quantified by the luciferase signal produced as a result of NFAT pathway activation. Luminescence was measured after 22 hours of incubation at 37°C with 5% CO_2_ with a luminometer using the Bio-Glo-TM Luciferase Assay Reagent according to the manufacturer’s instructions (Promega).

ADCC assays were performed using SARS-CoV2 CHO-K1 cells (genetically engineered to stably express a HaloTag-HiBit-tagged) as target cells and PBMC as effector cells at a E:T ratio of 30:1. HiBit-cells were seeded at 3,000 cells/well and incubated for 16 hours at 37°C, while PBMCs isolated from fresh blood were cultivated overnight at 37°C 5% CO2 in the presence of 5 ng/ml of IL-2. The day after, media was removed from the target cells and titrated concentrations of mAbs were added before the addition of PBMCs at 100,000 cells/well. As 100% specific lysis, Digitonin at 100 ug/ml was used. After 4 hours of incubation at 37°C, ADCC was measured with Nano-Glo HiBiT Extracellular Detection System (Promega; Cat. Nr.: N2421) using a luminometer (Integration Time 00:30). The assays were performed on 3 different donors and AUC (Area Under the Curves) were calculated.

ADCP assay was performed using ExpiCHO cells stably expressing full-length wild-type SARS-CoV-2 S (target cells) and labelled with PKH67 (Sigma Aldrich) as targets. PMBCs from healthy donor were labelled with CellTrace Violet (Invitrogen) and used as source of phagocytic effector cells. Target cells (10,000/well) were incubated with titrated concentrations of mAbs for 10 min and then mixed with PBMCs (200,000/well). The next day, cells were stained with APC-labelled anti-CD14 mAb (BD Pharmingen), BV605-labelled anti-CD16 mAb (Biolegend), BV711-labelled anti-CD19 mAb (Biolegend), PerCP/Cy5.5-labelled anti-CD3 mAb (Biolegend), APC/Cy7-labelled anti-CD56 mAb (Biolegend) for the identification of CD14+ monocytes. After 20 minutes, cells were washed and fixed with 4% paraformaldehyde before acquisition on a ZE5 Cell Analyzer (Biorad). Data were analyzed using FlowJo software. The % ADCP was calculated as % of monocytes (CD3- CD19- CD14+ cells) positive for PKH67. The assays were performed on 2 different donors and AUC (Area Under the Curves) were calculated.

### Hamster challenge experiment

Animal experiments were carried out by two independent laboratories. For the results shown in Fig.4 (A-C), 64 male golden Syrian hamsters (*Mesocricetus auratus;* RjHan:AURA) of 5-6 weeks of age (average weight 60-80 grams) were purchased from Janvier Laboratories (Le Genest-Saint-Isle, France) and handled under specific pathogen-free conditions. The animals were housed and manipulated in isolators in a Biosafety level-3 facility, with *ad libitum* access to water and food. Before manipulation, animals underwent an acclimation period of one week. Twenty-four hours before infection, the hamsters received an intraperitoneal injection of different concentrations of the hamsterized monoclonal antibodies (mAb) S309 (0.6, 1.7, 5 and 15 mg/kg), S2X324 (0.2, 0.6, 1.7 and 5 mg/kg) or the control isotype MGH2 (15 mg/kg). Hamsterization of human antibodies was based on the use of human VH and VL regions fused with the heavy and light chain constant regions of *Mesocricetus auratus IgG2a:* CH:ATTTAPSVYPLAPGGTPDSTTVTLGCLVKGYFPEPVTVSWNSGALTSGVHTFPSVLHSGLYSLSSSVTVPSSTWPSQTVTCNVAHPASSTKVDKKIEPRSCTSLPTLCPKCPAPDLLGGPSVFIFPPNPKDVLTISLTPKVTCVVVDVSEDEPDVQFNWFVNNVEVKTAETQPRQQQFNSTYRVVSSLPIQHQDWLSSKEFKCKVNNKALPSPIEKTISKPRGQARIPQVYTLPPPTEQMTQKVVSLTCMITGFFPADVHVEWEKNGQPEQNYKNTSPVLDTDGSYFMYSKLNVPKSSWEQGNIYVCSVLHEALRNHHTTKAISRSLGN CK:RSDAKPTVSIFPPSSEQLQSGSASLVCFVNNFYPKDINVKWKVDGSEKRDGVLQSITDQDSKDSTYSLSSTLTLTKGDYDSHNLYACEVTHKTSSTPIVKSLNKNEC CL:GQPSAAPSVTLFPPSSEELKTNQATLVCMIKEFYPSDVKVTWESDGIPITQGVKTTQPSKRDNKYLATSFLTMTAEAWKSRNSISCQVTHGGTTVEKSLSPAACF

Animal infection was performed as previously described (*95*). Briefly, the animals were anesthetized with an intraperitoneal injection of 200 mg/kg ketamine (Imalgène 1000, Merial) and 10 mg/kg xylazine (Rompun, Bayer), and 100 µL of physiological solution containing 6×10^4^ PFU of SARS-CoV-2/VoC delta (GISAID ID : EPI_ISL_2029113, supplied by Dr Olivier Schwartz, Institut Pasteur, Paris, France) was then administered intranasally to each animal (50 µL/nostril). Mock-infected animals received the physiological solution only. Infected and mock-infected hamsters were housed in separated isolators and were followed-up daily, for four days, when the body weight and the clinical score were noted. At day 4 post-inoculation, the animals were euthanized with an excess of anesthetics (ketamine and xylazine) and exsanguination. Blood samples were collected by cardiac puncture; after coagulation, the tubes were centrifuged at 2,000 x g during 10 min at 4°C, the serum was collected and frozen at -80°C until further analyses. The lungs were collected, weighted and frozen at -80°C until further analyses.

For the results shown in Fig.4 (D-E), the hamster infection model of SARS-CoV-2 including the associated analytical procedures, have been described before (*96, 97*). Briefly, female Syrian hamsters (Mesocricetus auratus) of 6-8 weeks old were anesthetized with ketamine/xylazine/atropine and inoculated intranasally with 50 μL containing 1×10^4^ TCID_50_ Delta (B.1.617.2; EPI_ISL_2425097) or BA.2 (*98*). Animals were treated once by intraperitoneal injection 24h before or after SARS-CoV-2 challenge (i.e., prophylactic vs therapeutic administration) with hamsterized S2X324 mAb. Hamsters were monitored for appearance, behavior and weight. At day 4 pi, hamsters were euthanized by i.p. injection of 500 μL Dolethal (200mg/ml sodium pentobarbital, Vétoquinol SA). Lungs were collected for viral RNA and infectious virus quantification by RT-qPCR and end-point virus titration, respectively. Serum samples were collected at day 4 pi for analysis of Ab levels. Animals with circulating Ab levels below the detection limit, indicating misdosing, were excluded.

### SARS-CoV-2 RT-qPCR

For data shown in Fig.4C, frozen lungs fragments were weighted and homogenized with 1 mL of ice-cold DMEM (31966021, Gibco) supplemented with 1% penicillin/streptomycin (15140148, Thermo Fisher) in Lysing Matrix M 2 mL tubes (116923050-CF, MP Biomedicals) using the FastPrep-24™ system (MP Biomedicals), and the following scheme: homogenization at 4.0 m/s during 20 sec, incubation at 4°C during 2 min, and new homogenization at 4.0 m/s during 20 sec. The tubes were centrifuged at 10.000 x g during 1 min at 4°C. Afterwards, 125 µL of the tissue homogenate supernatant were mixed with 375 µL of Trizol LS (10296028, Invitrogen) and the total RNA was extracted using the Direct-zol RNA MiniPrep Kit (R2052, Zymo Research). The presence of SARS-CoV-2 RNA in these samples was evaluated by one-step RT-qPCR in a final volume of 12.5 μL per reaction in 384-wells PCR plates using a thermocycler (QuantStudio 6 Flex, Applied Biosystems). Briefly, 2.5 μL of RNA were added to 10 μL of a master mix containing 6.25 μL of 2X reaction mix, 0.2 µL of MgSO_4_ (50 mM), 0.5 µL of Superscript III RT/Platinum Taq Mix (2 UI/µL) and 3.05 μL of nuclease-free water containing the nCoV_IP2 primers (nCoV_IP2-12669Fw: 5’-ATGAGCTTAGTCCTGTTG-3’; nCoV_IP2-12759Rv: 5’- CTCCCTTTGTTGTGTTGT-3’) at a final concentration of 400 nM, and the nCoV_IP2 probe (5’-FAM-AGATGTCTTGTGCTGCCGGTA-3’-TAMRA) at a final concentration of 200 nM (*99*).The amplification conditions were as follows: 55°C for 20 min, 95°C for 3 min, 50 cycles of 95°C for 15Ls and 58°C for 30 s, and a last step of 40°C for 30 s. Viral load quantification (expressed as RNA copy number/g of tissue) was assessed by linear regression using a standard curve of six known quantities of RNA transcripts containing the *RdRp* sequence (ranging from 10^7^ to 10^2^ copies).

For data shown in Fig. 4 D, E, hamster lung and trachea tissues were collected after sacrifice and were homogenized using bead disruption (Precellys) in 350 µL TRK lysis buffer (E.Z.N.A.® Total RNA Kit, Omega Bio-tek) and centrifuged (10.000 rpm, 5 min) to pellet the cell debris. RNA was extracted according to the manufacturer’s instructions. RT-qPCR was performed on a LightCycler96 platform (Roche) using the iTaq Universal Probes One-Step RT-qPCR kit (BioRad) with N2 primers and probes targeting the nucleocapsid (*96*). Standards of SARS-CoV-2 cDNA (IDT) were used to express viral genome copies per mg tissue.

### End-point virus titrations

Lung tissues were homogenized using bead disruption (Precellys) in 350 µL minimal essential medium and centrifuged (10,000 rpm, 5min, 4°C) to pellet the cell debris. To quantify infectious SARS-CoV-2 particles, endpoint titrations were performed on confluent Vero E6 cells in 96-well plates. Viral titers were calculated by the Reed and Muench method (*100*) using the Lindenbach calculator and were expressed as 50% tissue culture infectious dose (TCID_50_) per mg tissue.

### MD simulations

Structures of the Wuhan-Hu-1 and BA.1 RBDs were prepared as previously described (*101, 102*). The coordinates of the Wuhan-Hu-1 RBD were obtained from PDB 6M0J (*44*) for which ACE2 was removed and as previously described (*26*) the glycan at position 343 added in the RBD using ISOLDE (*103*) to visually place, link and minimize each monosaccharide beyond the *N*-acetylglucosamine for which there was density. The resulting model was then prepared using QuickPrep (MOE v2020.0901, https://www.chemcomp.com). The coordinates of the BA.1 RBD were obtained from PDB 7TN0 (*17*) with the complex glycan at position 343 added before refinement using ISOLDE and model preparation using QuickPrep. The coordinates of the BA.2 RBD were generated by mutating positions 371, 376, 405, and 408 of the refined BA.1 structure, optimizing the rotamers at the four positions using Repack in Protein Builder (MOE v2020.0901), followed by model preparation using QuickPrep.

The 3 RBD structures were each parameterized using tleap with the Amber ff14SB force field (*104*) for the protein, GLYCAM_06j-1 for the glycan (*105*), TIP3P for water (truncated octahedral cell with 18 Å buffer around solute) (*106*); the Joung & Cheatham parameters (*107*) were used for the ions neutralizing the charged solute (2 Cl- ions for Wuhan-Hu-1, 6 Cl- ions for BA.1 and BA.2) and the additional 0.15 M excess Na+ and Cl- ions (80 Na+ and 80 Cl- for Wuhan-Hu-1, 76 of each for BA.1 and BA.2).

We generated 4.8 μs of aggregate MD for each of the three RBDs by performing eight independent MD simulations seeded with different initial velocities. Each of these 24 simulations was subjected to a nine-stage minimization and equilibration procedure as previously described (*101*). This involved 10,000 steps of restrained minimization with a 100 kcal/mol Å^2^ force constant applied to all heavy atoms resolved in the structure followed by 100 ps of restrained MD heating to 300 K, 100 ps of restrained MD at 300 K, 250 ps of restrained MD with a 10-fold reduction in restraint force constant at 300 K, 10,000 steps of backbone-restrained minimization, 100 ps of backbone-restrained MD, 100 ps of backbone-restrained MD at 300 K, 100 ps of backbone-restrained MD with a further 10-fold reduction in restraint force constant, 100 ps of backbone-restrained MD with another 10-fold reduction in restraint force constant (0.1 kcal/mol Å^2^), and 2.5 ns of unrestrained MD at 300 K. An additional 0.6 μs of unrestrained MD simulation was performed for each of the 24 independent simulations.

Simulations were processed using cpptraj (*108*) to image the coordinates, RMS-align RBD CL atoms to the S309:RBD complex (PDB ID: 7TN0), and to strip solvent and ions. Volumetric density maps were computed using cpptraj with grid points spaced 0.25 Å over a grid with dimensions 279 Å, 296 Å, 307 Å. This resulted in Cartesian coordinates of grid points and the probability of finding any glycan atom at those grid points. This procedure was repeated for S309 from the S309:RBD complex used to align RBDs above. The maximum probability (isovalue) at which no glycan atom from RBD Wuhan-Hu-1 occupied a grid point also occupied by S309 was 0.057 and this isovalue was used for subsequent analysis with BA.1 and BA.2. Overlap was calculated by counting the grid points occupied by S309 (probabilities were one or zero because it was a static structure) and any glycan atom if the probability for the glycan was 0.057 or less at that grid point. Smaller isovalues increase the volume size because ephemeral glycan conformations are included in the map whereas a larger isovalue (occupancy probability) reduces the volume size because only persistent glycan conformations are included in the map. The total volume housed by a volumetric density map for a glycan was computed by counting the grid points occupied by the glycan with an isovalue of 0.057. Renderings for volumetric density maps of the glycan were generated using VMD (*109*).

